# A Mechanical Theory for the Formation of Short Association Fibers in the Brain

**DOI:** 10.64898/2026.08.12.744475

**Authors:** Akbar Solhtalab, Jixin Hou, Guannan Cao, Zhibin He, Lana Pierotich, Tuo Zhang, Kara E. Garcia, Xianqiao Wang, Mir Jalil Razavi

## Abstract

The development of neural connections in the brain results from a complex interplay between biological processes and mechanical forces. A key question in neuroscience is how physical forces and the mechanical properties of brain tissue influence the formation of structural connections. Here, we demonstrate that mechanical forces play an essential role in shaping the emergence of short-range connections, particularly U-shaped fibers that link neighboring regions of the cortex. Using a computational model that incorporates our “stress-dependent axon reorientation” hypothesis, we simulate how growing axons respond to the mechanical stress field generated by cortical folding. Our results suggest that axonal growth and reorientation may be strongly influenced by local mechanical cues, helping establish the organization of these short-range pathways. Supported by *in vivo* diffusion tensor imaging and histological observations, our findings provide a physical explanation for why these fibers predominantly adopt U-shaped trajectories, and why connections between gyri (ridges) are more prevalent than those between sulci (valleys) or spanning gyri and sulci. These results suggest that understanding the mechanics of brain folding is critical for fully explaining the formation of brain connectivity and its variations in health and disorder.

**Teaser:** Mechanical forces during cortical folding guide the formation of short association fibers in the brain.

## Introduction

Short association fibers (SAFs) are short intrahemispheric cortico-cortical fibers that connect nearby cortical regions. Their most superficial and well-known component is the classical U-fibers, which arch beneath the cortex to connect adjacent gyri(*1*, *2*). Located in the superficial white matter just below the gray matter–white matter boundary, these fibers typically adopt U-shaped geometries that follow the cortical folding pattern. SAFs play a significant role in brain development(*3*, *4*), contribute to overall white matter volume, and are crucial for brain function(*5*, *6*). In addition, alterations in their structure and connectivity have been implicated in neurological disorders such as autism spectrum disorder (ASD)(*7*), epilepsy(*8*, *9*), and Alzheimer’s disease(*10*, *11*).

SAFs develop in parallel with cortical folding, beginning to grow during fetal development and continuing to grow throughout infancy before ultimately undergoing myelination(*12–14*), with maturation extending into childhood and adulthood(*15*). Despite their recognized importance, the developmental mechanisms governing SAFs formation remain largely unexplored. Notably, U-fibers, which are a major subset of SAFs, emerge later than deeper white matter tracts(*16–18*), suggesting that their growth may be regulated by distinct biological processes. Their spatial organization, which closely follows cortical folding patterns, points to a potential link between the cortical folding process and the arrangement of these fibers. Figure 1 illustrates SAFs, including U-fibers, in the human brain, shown through both anatomical dissection (Fig. 1A) and fiber tractography derived from diffusion tensor imaging (DTI) data (Fig. 1B). However, SAFs remain underrepresented in structural connectivity studies, in part due to the inherent challenges of mapping these short-range fibers using DTI. Their close proximity to the cortical surface, sharp curvature, and complex trajectories often limit the reliability of DTI-based reconstructions. Ongoing research continues to advance the robustness of mapping SAFs(*1*, *19*, *20*), aiming to overcome these limitations and improve the accuracy of detecting and quantifying their architecture.

**Fig. 1.**
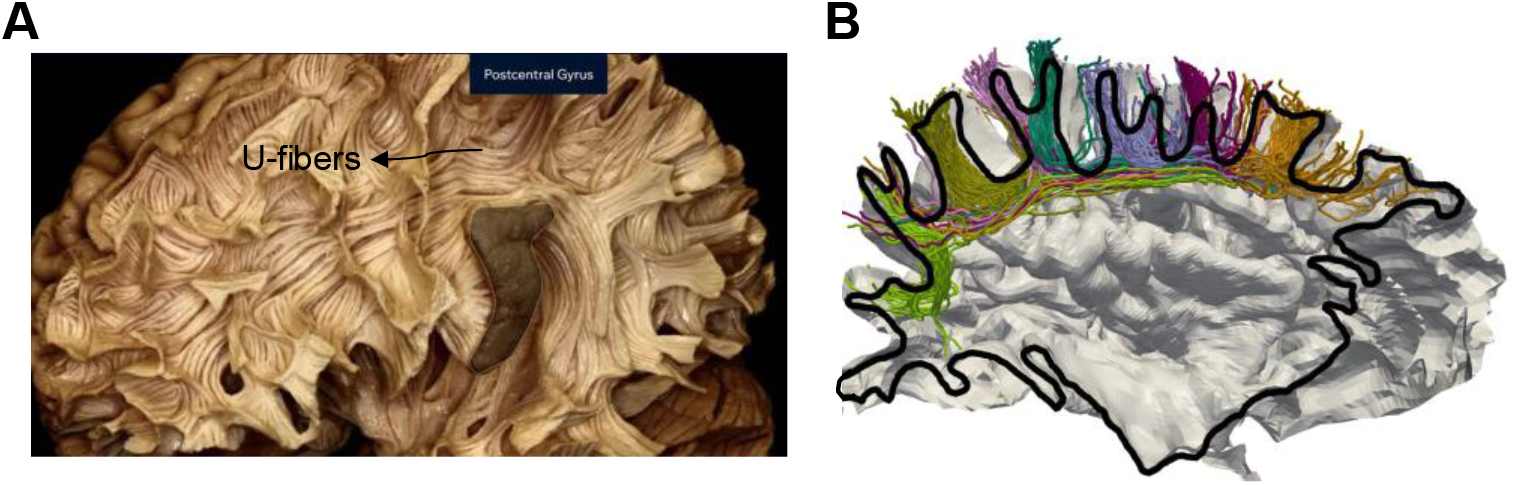
Depiction of short association fibers (SAFs) in the human brain. **(A)** A dissection image shows the physical bundles of short association fibers. U-fibers are visible coursing beneath sulci. This image was adapted and used with permission from the Neurosurgical Atlas by Aaron Cohen-Gadol, MD. **(B)** Joint representation of the cortical surface (boundary between white and gray matter, derived from structural magnetic resonance imaging (sMRI)) and association fiber streamlines (colored curves, derived from diffusion magnetic resonance imaging (dMRI)) on a sagittal section (outlined in black) of a mature brain from the Human Connectome Project (young adult) dataset. Dominant fiber bundles within each gyrus were manually separated and color-coded for improved visualization. SAFs predominantly form U-shaped patterns, connecting adjacent gyri and coursing along the sulcal fundi. Figures (A) and (B) have not been generated from the same brain.

There has been growing interest in the role of mechanics in brain folding and connectivity development(*21–32*), particularly in relation to two longstanding theories: the differential growth theory proposed by Richman et al.(*33*) and the axonal tension theory by Van Essen(*34*, *35*). The differential tangential growth theory posits that faster expansion of the outer cortical plate compared to the inner subplate creates mechanical instabilities that drive the formation of gyri (ridges) and sulci (valleys). Meanwhile, the axonal tension theory proposes that tension generated by axonal connections influences brain folding by pulling interconnected regions together. While the differential tangential growth theory is widely supported by physical models(*28*, *36*, *37*), recent evidence suggests that axonal tension does not induce cortical folding, although axons are under significant tension(*29*, *38*). Instead, cortical folding appears to exert mechanical forces on axons, stimulating their elongation and promoting white matter growth(*22*, *39*).

Recently, simulations based on differential growth theory have evolved to recognize the importance of axons within the white matter. Some simulations have begun to consider white matter, composed primarily of axonal fibers, as a material that undergoes growth (axon elongation) in response to mechanical stretch(*23*, *40*, *41*). While axonal tension may not directly induce folding, it could play a key role in regulating folding patterns and ensuring consistent fold placement(*29*, *42*, *43*). Moreover, the patterns of stress induced by folding are sufficient to induce remodeling of axonal fibers consistent with the formation of short and long association fibers(*22*). Despite these efforts to model the effect of axonal fibers in the folding process, either using stretch-driven continuum models(*22*, *23*, *29*, *40*) or explicit but passive fibers(*29*, *42–44*), the mechanics of cortical folding and connectivity development is not well understood to date, particularly concerning the mechanics of emerging SAFs in the developing brain.

In this study, we investigate the biomechanical mechanisms guiding the formation and organization of SAFs, particularly U-fibers, during brain development. Building on our recent work(*45*), we hypothesize that the proposed “stress-dependent axon reorientation” mechanism, driven by the mechanical environment of cortical folding, plays a central role in shaping SAF trajectories. This mechanism suggests that, in addition to stress-dependent growth, axons reorient toward the direction of maximum principal tensile stress to establish stereotyped subcortical fiber pathways. To test this hypothesis, we develop a multiscale mechanical model to investigate the formation and dynamic pathfinding of both afferent and efferent fibers in a folding brain. Our simulations, incorporating stress-dependent axon growth and reorientation, offer a potential mechanical explanation for the formation and patterning of SAFs. The simulation results are compared with *in vivo* DTI data from the human brain and histological observations from other gyrencephalic species, providing insights into the mechanical processes that shape SAFs geometries. In particular, we examine why gyrus-gyrus SAF connections are more prevalent than gyrus-sulcus or sulcus-sulcus. In addition, we explore how the timing of fiber initiation relative to cortical folding, axonal growth rate, and the mechanical properties of brain tissue influence SAF patterning. While recognizing that connectivity patterning is shaped by a complex and precise orchestration of molecular, cellular, and mechanical processes(*46*), this study represents a significant step toward understanding the mechanistic underpinnings of SAF development in both healthy and pathological conditions.

## Results

To shed light on the dynamic interaction between folding and connectivity development, we investigated how afferent and efferent axonal fiber bundles (hereafter referred to as fibers) grow and navigate within a system undergoing cortical folding according to the proposed stress-dependent axon reorientation mechanism. In this study, SAFs refer to a specific subset of cortico-cortical fibers that primarily connect adjacent gyri, often forming U-shaped trajectories beneath the cortical surface (U-fibers). Additionally, SAFs may extend to connect closely situated gyri within the same hemisphere. These fibers are anatomically distinct from long association fibers, such as the superior longitudinal fasciculus, which span greater distances to connect cortical regions across different lobes. Results derived from the models and imaging data are presented here to investigate potential mechanisms relevant to brain development, rather than directly replicating the precise processes occurring in the human brain.

### Axonal pathfinding within a stress field

Figure 2 illustrates the dynamic initiation and development of both afferent and efferent fibers within the stress field of the subplate (later white matter) during cortical expansion and folding. In this study, the term subplate refers to the non-fiber components of subcortical tissue, including the extracellular matrix (ECM) and cellular components (see Methods). The simulation time was defined as model time *t*, ranging from 0 to 50 days, and was normalized using the dimensionless simulation time *T* = *G^c^t*, where *G^c^* is the cortical areal growth rate parameter. In all simulations, *G^c^* = 0.02 *d*^−1^; therefore, *T* = 0 corresponds to the beginning of cortical expansion and *T* = 1 corresponds to the final folded configuration after 50 model days. Here, *t* denotes the model time in days, and *T* provides a normalized measure of developmental progression. Parameters reported in units of days, including the axonal growth rate and stress-dependent elongation coefficient, are evaluated with respect to this model time *t*. The gestational-week windows discussed later are used as biologically informed staging labels for relative developmental timing, rather than as a direct conversion from model days to fetal age.

**Fig. 2.**
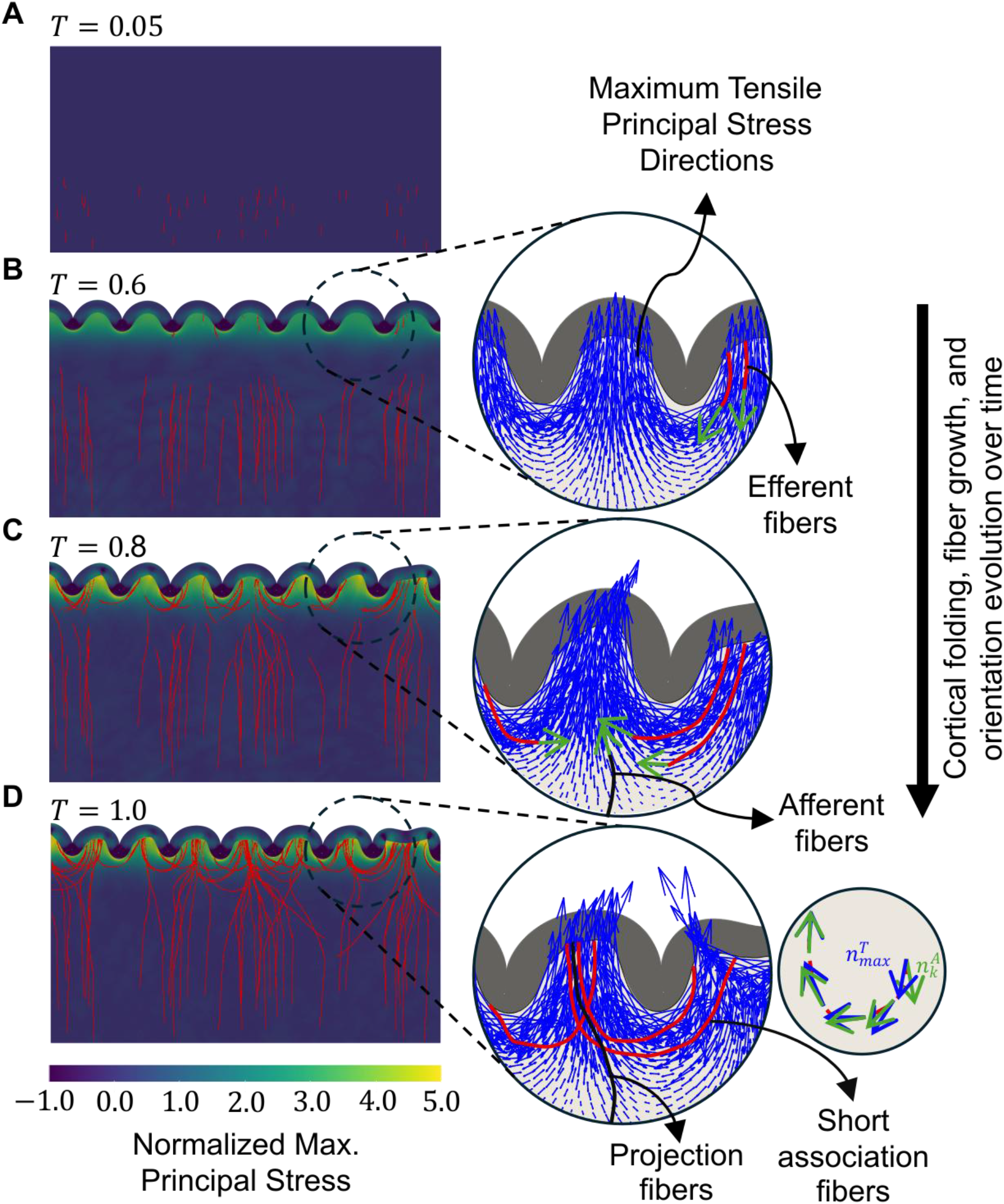
Stress field of the folding bilayer system alters the growth trajectories of navigating afferent and efferent fibers. (**A**) Afferent fibers originate from deep brain layers and extend toward the cortical plate before folding begins. (**B**) After folding initiates, efferent fibers, representing prospective short association fibers (SAFs), grow from the cortex–subplate interface into the subplate. As folding progresses, the maximum tensile principal stress (MTPS) directions (blue arrows) become predominantly horizontal beneath sulci and more vertical beneath gyri, causing efferent fibers to gradually reorient toward the local MTPS. (**C**) Afferent fibers approach high-stress regions near the cortex and also reorient toward the evolving MTPS field, while efferent fibers continue to grow and adapt to the stress environment. (**D**) At the final stage, afferent fibers reach the cortical plate, whereas efferent fibers form SAF-like trajectories connecting adjacent cortical regions. The simulation time *T* = *G*^c^*t* is dimensionless, where *G*^c^ represents the cortical areal growth rate parameter and *t* is the elapsed time. *T* ranges from 0 to 1 to capture the progression of cortical folding development. The normalized principal stress is calculated as the ratio of the maximum principal stress to the shear modulus of the subplate at each simulation step. A single, consistent color scale is used across all panels to allow direct comparison of normalized stress values. A reduced set of 100 fibers (out of 150) is used solely for visualization purposes to improve clarity in illustrating fiber trajectories and their interaction with the evolving cortical geometry. The used parameters in this figure are: *μ*_f_/*μ*_s_ = 2, *μ*_c_/*μ*_s_ = 2, *a* = 0.015 mm Pa^−1^ d^−1^, *G*^axn^ = 0.6 mm d^−1^, where *μ*_f_, *μ*_s_, and *μ*_c_ are the shear moduli of the fiber, subplate, and cortex, respectively, *a* is the stress-dependent elongation rate, and *G*^axn^ is the baseline axon growth rate.

From *T* = 0 to *T* = 1 afferent and efferent fibers initiate, elongate, navigate, and become established in the stress landscape of the folding system. In Fig. 2, we used a baseline axonal growth rate of *G*^axn^ = 0.6 mm d^−1^, a stress-induced elongation rate of *a* = 0.015 mm Pa^−1^ d^−1^, and a relaxation parameter of *t*^∗^ = 200. These parameters are defined with respect to physical time, while pseudo-time *T* is used only to represent the normalized progression of the system. *G*^axn^ is a model parameter that represents the daily axonal growth rate in the absence of external forces, *a* quantifies the additional axonal elongation in response to external tensile stresses, and *t*^∗^ controls the reorientation rate (see Methods). Biologically, *t*^∗^ represents an effective mechanobiological response timescale that captures the delayed reorientation of axons to anisotropic mechanical cues. It reflects the combined duration of mechanosensing through adhesion complexes and mechanosensitive ion channels, intracellular signaling such as calcium influx, and subsequent cytoskeletal and traction force remodeling that redirect growth cone motion(*47*, *48*). In this context, *t*^∗^ governs how rapidly axons can realign with the local mechanical field, such as the maximum principal tensile stress (MTPS). Alignment analysis (angle between fiber orientation and MTPS) reveals a gradual, time-dependent response, indicating that fibers progressively reorient toward the MTPS direction under the influence of *t*^∗^ within the evolving stress field (Fig. S1).

Early in cortical expansion, almost no tension exists at the deep part of the subplate, allowing afferent fibers to grow in more randomized paths, as illustrated in Fig. 2A-B. Initial angles for these fibers are randomly assigned within a range of *π*/2 ± 10% to direct them towards the cortical plate similar to the real brain. In this specific case, efferent fibers (fibers originating at the cortex-subplate boundary) begin to grow after folding onset, with initial angles in the range 3*π*/2 ± 10%, guiding them from the cortical plate toward deep brain structures.

Primary folding in the fetal brain is thought to be guided by genetically patterned, regionally heterogeneous radial growth and progenitor activity, which establish stereotyped differences in cortical expansion and initiate the formation of primary gyri and sulci(*49*). Following the establishment of primary folds, the differential tangential growth hypothesis posits that rapid cortical expansion relative to the slower growing subplate generates compressive stresses within the cortical layer, driving its buckling into the subplate and leading to the formation of secondary and tertiary folds. Blue arrows in the zoomed views in Fig. 2 represent the directions of MTPS in the subplate, while green arrows indicate the preferred growth direction at the fiber tip. As cortical folding progresses, MTPS intensifies, with predominantly tangential tension beneath emerging sulci and primarily radial tension beneath gyri, as depicted in Fig. 2B. Figure 2C-D reveal that as MTPS increases, a stress-dependent reorientation of fiber’s growth trajectory occurs, with preferred directions (green arrows) gradually aligning with their respective MTPS directions (blue arrows). By the completion of folding at *T* = 1, the reorientation aligns nearly all fibers with MTPS directions (Fig. 2D). These results are consistent with previous predictions of tension-induced fiber remodeling in a continuum framework(*22*).

Figure 2 also depicts the normalized principal stress at each simulation step, defined as the ratio of the maximum principal stress to the shear modulus of the subplate, to illustrate the evolving stress landscape within the subplate and its influence on fiber pathfinding during cortical folding. Efferent fibers that initially grow toward the core of the model progressively respond to the folding induced stress field, which reorients their trajectories back toward the cortical plate. This process ultimately leads to the formation of SAFs, which predominantly adopt U-shaped paths. However, some efferent fibers escape high-stress regions without reorienting into U-shaped trajectories and instead form non-SAFs (projection-type fibers) that extend from the cortical plate into the deeper subplate. This behavior arises from the interplay between initial fiber orientation, local stress-field structure, and finite reorientation dynamics. Fibers that are initially aligned with the MTPS direction can rapidly traverse high-stress regions before significant reorientation occurs, while others fail to adapt due to the rapid evolution of the stress field near bifurcation. Similarly, afferent fibers navigating toward the cortical plate encounter the stress field, which may bias their trajectory more toward gyri than sulci^54^. In the baseline case shown in Fig. 2, efferent fibers are initiated after the onset of folding to isolate the influence of an already-developed stress field on axonal pathfinding. However, during brain development, axonal growth occurs over a broad time window, and efferent fibers can also emerge earlier, prior to the onset of cortical folding. Because this timing strongly influences connectivity patterns, the dependence of SAF development on its temporal association with cortical folding is examined in detail in the following section.

### Effect of timed fiber emergence on SAFs patterning

Neurodevelopmental studies have shown that early-emerging cortical efferents, which typically arise from deep-layer neurons born early in development, tend to establish long-range connections before the onset of major folding events, whereas late-emerging efferents from upper-layer neurons tend to form more localized or short-range connections as the cortex matures and folds(*50*, *51*). Therefore, we investigated how the timing of fiber-growth initiation relative to the onset of cortical folding influences the organization and connectivity of SAFs. For better representation of the results, we suppressed the development of the afferent fibers in the remaining models of this study. However, Fig. S2 shows that the inclusion of afferent fibers has no effect on the behavior of SAFs and their developmental characteristics.

To investigate how the relative timing of fiber emergence influences SAF formation, we grouped the simulations into four timing regimes defined with respect to the bifurcation time *T_β_*. Here *T_β_* is defined as the onset time of mechanical instability associated with cortical folding. Specifically, it corresponds to the point at which the initially flat cortical surface becomes unstable under compressive stresses, leading to the initiation of buckling and the emergence of surface undulations. This time marks the transition from a pre-folding stable state to a folding regime; it does not correspond to a fully developed fold or a specific amplitude threshold.

Categorizing the onset of fiber growth (*T_init_*) relative to the bifurcation time, provides an opportunity to understand how this temporal association impacts fiber trajectories, cortical folding, and connectivity. A biologically informed interpretation of these regimes, including characteristic SAF morphology at each stage, is provided in Table S1. These regimes are defined to distinguish whether fibers emerge before, near, during, or after the establishment of the folding-induced stress field, rather than to represent sharply defined biological phases. Accordingly, the fractional cutoffs are modeling thresholds used to test the qualitative ordering of fiber outcomes, from projection-dominant early emergence to U-fiber-dominant late emergence, rather than biologically calibrated boundaries or exact gestational transitions.

Categorizing the onset of fiber growth, *T_init_*, relative to *T_β_* allows us to examine how the sequence of fiber emergence and cortical folding affects fiber trajectories and connectivity. A qualitative comparison between the model-derived ordering of SAF morphologies and the developmental ordering reported in fetal observations is provided in Table S1. These regimes are defined to distinguish whether fibers emerge before, near, during, or after the establishment of the folding-induced stress field, rather than representing sharply defined biological phases. Accordingly, the fractional cutoffs are modeling thresholds used to test the qualitative ordering of fiber outcomes—from projection-dominant early emergence to U-fiber-dominant late emergence—rather than biologically calibrated boundaries or exact gestational transitions.

The four computational timing regimes are defined as follows. For each regime below, the model-predicted SAF morphology is compared qualitatively with fetal observations; the reported gestational intervals are not temporally mapped to the computational regimes.

1. Pre-folding regime 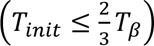: Fibers that initiate growth well before the bifurcation process tend to avoid the high-stress regions generated by cortical folding. As shown in Fig. 3A, these fibers predominantly form descending long projection trajectories, establishing connections between cortical regions and deep brain structures. Their growth direction is largely influenced by intrinsic stochastic tip dynamics, which guide them downward before encountering the folding-induced high-stress regions with U-shaped MTPS directions. Under these conditions, most fibers develop into long projection fibers, while a smaller subset forms U-fibers. As a qualitative biological comparison, fetal imaging at approximately 22–26 GW shows relatively flat, loosely organized, and multidirectional superficial association pathways with limited influence from folding(*52*). The common morphological feature is the limited folding-induced organization and absence of predominantly well-defined U-shaped bundles.
2. Peri-folding onset regime 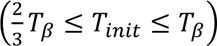: Fibers that begin growth closer to the bifurcation time are more likely to traverse regions that are starting to experience stress gradients even before they are fully folded. These fibers exhibit mixed behaviors, forming both projection fibers and early-stage U-fibers. The stress distribution during this phase starts to influence fiber trajectories, as shown in Fig. 3B. A comparable morphological transition has been reported at approximately 26–30 GW, when fetal SAFs begin to exhibit partial organization alongside emerging cortical folds(*52*). In both cases, weakly organized trajectories progressively give way to more curved and localized pathways under the increasing influence of folding.
3. Active-folding regime 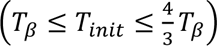: Fibers initiating growth during or shortly after bifurcation are strongly influenced by the emerging folds and associated high-stress regions. As shown in Fig. 3C, these fibers are more likely to reorient toward the MTPS direction, forming U-fibers that connect adjacent cortical regions. The folding process acts as a scaffold, directing the fibers toward localized paths within the developing gyri and sulci. Consistent with this qualitative progression, fetal observations at approximately 30–34 GW show substantial SAF reorientation and an increasing prevalence of curved trajectories associated with developing cortical geometry(*52*). The shared feature is pronounced folding-related reorientation and the emergence of localized U-shaped connections.
4. Late-folding regime 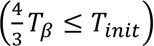: Late-growing fibers encounter fully developed cortical folds and an established stress landscape. These fibers primarily contribute to short, localized connections within the same gyrus or between neighboring regions. The advanced stage of cortical folding restricts their trajectories, confining them to high-stress regions that favor the formation of U-fibers (Fig. 3D). Fetal imaging at approximately 34–38 GW and later similarly demonstrates increasingly organized U-shaped bundles between neighboring gyri(*52*). The relevant similarity is the predominance of localized U-shaped organization once the folding geometry is substantially established. This category also applies to fibers that develop after cortical folding is complete (*T* > 1). In this case, fibers would be expected to follow the pattern of residual stress in the white matter(*53*), in the absence of significant stress relaxation, which may represent SAFs that primarily develop postnatally.

**Fig. 3.**
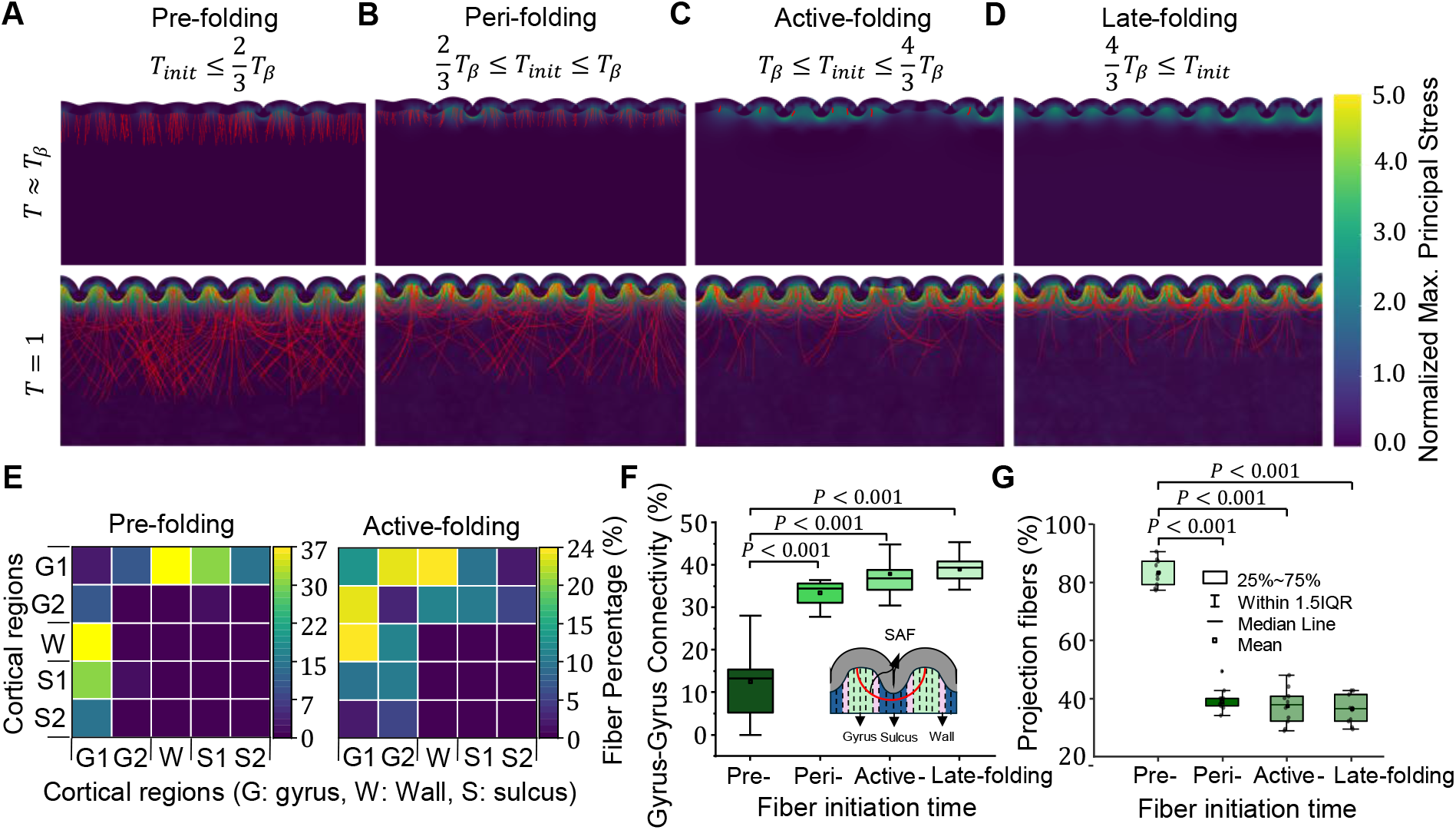
The association time between folding and connectivity development regulates the formation and patterning of short association fibers (SAFs). (**A**) Fibers start to grow before cortical folding (pre-folding growth). (**B**) Fibers start to grow close to cortical folding (peri-folding growth). (**C**) Fibers start to grow during cortical folding (active-folding growth). (**D**) Fibers start to grow after cortical folding (late-folding growth). (**E**) Connectivity distribution between cortical regions for two selected categories: pre-folding growth and active-folding growth. Rows and columns correspond to the same regions (G1, G2: gyri; W: wall; S1, S2: sulci), and the symmetric matrix represents pairwise connectivity. These regions are defined by dividing the cortical plate into five equal-area bins from sulci to gyri, as illustrated in Fig. 6D. (**F**) Gyrus-to-gyrus connectivity density across all categories, showing a significant increase in localized connectivity as fiber growth shifts to later stages. (**G**) Boxplots show the percentage of fibers that do not reorient toward MTPS directions for four initiation categories relative to the bifurcation time *T_β_*: pre-folding, peri-folding, active-folding, and late-folding. The results show a significantly higher frequency of projection-like fibers for early initiation compared to later initiation regimes (p<0.001). Statistical analyses were two-sided and used Tukey’s post hoc method for multiple comparisons. Box plots show the interquartile range (IQR; 25th to 75th percentile), with the mean shown as a white dot and the median as a black line. Whiskers indicate the data range within 1.5 times the IQR. The parameters used in this figure are as follows: *μ*_f_/*μ*_s_ = 2, *μ*_c_/*μ*_s_ = 2, *a* = 0.015 mm Pa^−1^ d^−1^, *G*^axn^ = 0.6 mm d^−1^, where *μ*_f_, *μ*_s_, and *μ*_c_ are the shear moduli of the fiber, subplate, and cortex, respectively, *a* is the stress-dependent elongation rate, and *G*^axn^ is the axon growth rate.

The connectivity distribution between different regions of the folded cortex varies considerably based on the timing of fiber growth initiation. For pre-folding, efferent fibers predominantly traverse long paths, often bypassing high-stress regions associated with cortical folding. This results in a more heterogeneous connectivity pattern, with fibers frequently forming connections between distant cortical regions, including both gyrus-to-gyrus and gyrus-to-sulcus pathways, as shown in Fig. 3A. In contrast, the active-folding phase demonstrates a predominant gyrus-to-adjacent-gyrus connectivity (Fig. 3C). The earlier onset of folding in this case creates localized high-stress regions that guide fibers to reorient and form SAFs within neighboring gyri. This trend is visually supported by the observed connectivity matrices, where active-folding cases exhibit dense and localized connections compared to the broader distributions seen in the pre-folding scenario (Fig. 3E).

Figure 3F illustrates the gyrus-to-gyrus connectivity across different categories of timed fiber emergence. In the pre-folding phase, the fiber density connecting gyri is comparatively low, as fibers tend to form long-range connections bypassing local cortical regions. However, as fibers initiate growth in later stages, the density of gyrus-to-gyrus connectivity increases significantly. Statistical analysis reveals a strong correlation between the pre-folding regime and subsequent regimes, underscoring the temporal dependence of fiber organization and patterning on cortical development dynamics. Despite the predominance of stress-dependent reorientation, a subset of fibers exhibits projection-like behavior, escaping high-stress regions without forming U-shaped trajectories (Fig. 3G). This behavior arises from the interplay between initial fiber orientation, local stress-field structure, and finite reorientation dynamics. Fibers that are initially aligned with the MTPS direction can rapidly traverse high-stress regions before significant reorientation occurs, while others fail to adapt due to the rapid evolution of the stress field near bifurcation. This effect is most pronounced for early initiation and decreases substantially for later stages as the stress field stabilizes.

### Effect of biomechanical parameters on SAF patterning

Understanding how biomechanical parameters influence the formation of SAFs is crucial, as these fibers play a key role in local connectivity and are implicated in various neurodevelopmental disorders. Changes in mechanical factors such as tissue stiffness and growth dynamics during development may disrupt the spatial patterning of SAFs, potentially leading to altered connectivity and functional deficits observed in conditions like ASD(*54*) and bipolar disorder(*55*).

We explored the influence of biomechanical parameters including axonal growth rate, cortical stiffness, and fiber stiffness on SAF patterning (Figs. S3-6). Cortical growth rate has been well characterized, and is known to vary over the course of development as well as cortical regions(*56*, *57*). By contrast, current estimates of axonal growth rate, cortical versus subcortical stiffness, and fiber versus subplate stiffness span a wide range of potential values(*58–60*). Notably, while simulations varying these biomechanical factors exhibit some differences, the key observations described in Figs. 2-3 remained consistent across all cases.

Simulations considering a higher axonal growth rate, or equivalently a reduced cortical plate growth rate, produced a lower SAF density while promoting the formation of longer projection fibers due to diminished stress-dependent reorientation in deeper subplate (Fig. S3). Increased axonal growth rates (or relative slowdown in cortical expansion) shift the timing of connectivity development relative to cortical folding. Similar to the previously observed effect of early fiber emergence, this shift alters the dominant connection patterns by reducing gyrus-to-gyrus connections and increasing gyrus-to-wall connectivity (Fig. 4A). A detailed discussion of the effects of the relative growth rates of the cortex and axons on SAF patterning is provided in the Supplementary Materials.

**Fig. 4.**
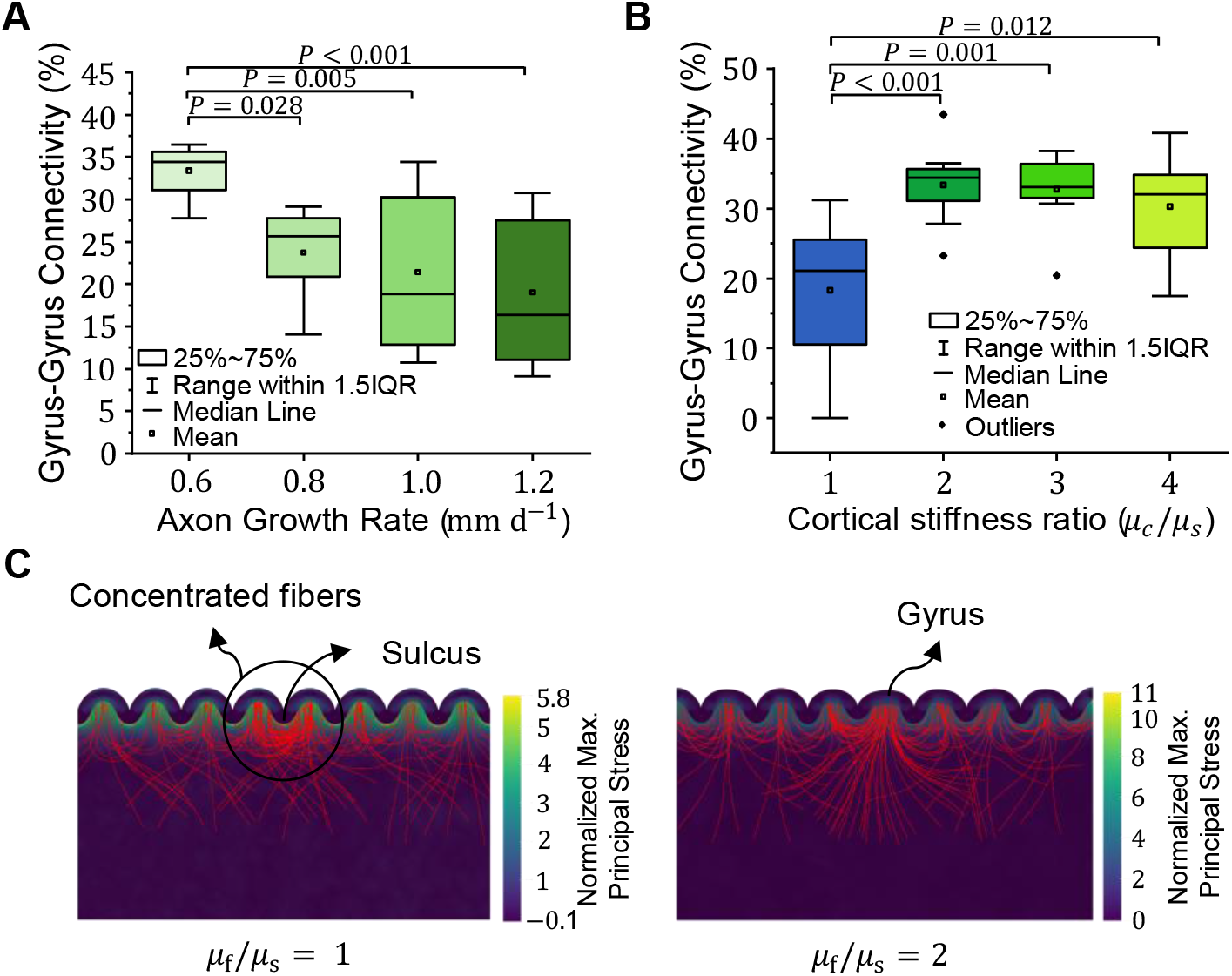
Biomechanical parameters control the patterning of short association fibers (SAFs). (**A**) Percentage of gyrus-to-gyrus connections as a function of fiber growth rate, showing a significant decrease with increasing growth rate. (**B**) Percentage of gyrus-to-gyrus connections for different cortex-to-subplate stiffness ratios, revealing a notable increase when the ratio exceeds 1. (**C**) Influence of fiber stiffness on the positioning of gyri and sulci. Left: *μ*_f_/*μ*_s_ = 1; Right: *μ*_f_/*μ*_s_ = 2, where *μ*_f_, *μ*_s_, and *μ*_c_ are the shear moduli of the fiber, subplate, and cortex, respectively. The box plots represent the interquartile range (IQR), extending from the 25th to the 75th percentile. The mean is depicted as a white dot, while the median is shown as a black line within the box. The whiskers indicate the data range within 1.5 times the IQR.

Simulation results also suggest that the stiffness ratio between the cortex and subplate influences the timing of folding, consistent with past studies(*25*). This effect is demonstrated in Fig. S4, where the shear modulus ratio of cortex to ECM is varied from 1 to 4. For a given window of fiber emergence, such shifts in folding timing directly influence the formation, connectivity, and organization of SAFs. Specifically, higher cortical stiffness accelerates folding, promoting earlier stress-dependent fiber reorientation and enhanced SAF formation, particularly between adjacent gyri (Fig. 4B). In contrast, lower stiffness delays folding, allowing more fibers to develop into long projection pathways. Importantly, Fig. S4 shows that, generally, the formation of SAFs remains robust across this range of material properties, indicating that the key findings do not depend on a specific parameter choice. A detailed discussion of the effects of the cortical to subplate stiffness ratio is provided in the Supplementary Materials.

Our findings also reveal that fiber stiffness affects fold localization and the structural organization of SAFs (Fig. 4C). When fiber and subplate stiffness are equal (*μ*_f_/*μ*_s_ = 1), fibers have no effect on fold positioning, and folding patterns remain symmetric and geometry-driven. However, when fibers are stiffer than the subplate (*μ*_f_/*μ*_s_ ≥ 2), regions with high fiber concentration consistently form gyri, suggesting that local fiber stiffness can direct fold localization (Fig. S5). While in this study we define fiber concentration as an initial condition that localizes fold formation, it is also possible that folding-induced stress fields could reciprocally influence fiber trajectories, potentially guiding fibers toward regions of radial tension, such as developing gyri(*45*). More detailed findings are presented in the Supplementary Materials, including the effect of fiber stiffness on fold placement throughout the full timeline of cortical folding and connectivity development, as shown in Fig. S6.

### Comparison of model predictions with experimental and imaging data

To further investigate the origin of SAF development and its relationship to cortical folding, we compared the modeling results with the experimental and imaging observations of gyrencephalic ferret and human brain (Fig. 5). Experimental evidence from ferret brains (Fig. 5A–B), adapted from Yoshino et al.(*16*), provides direct insight into the temporal emergence of U-fibers at the axonal level. Using in utero electroporation at embryonic day E31 followed by postnatal imaging at P0, P6, and P16, GFP-labeled axons show that fibers in the deep white matter are already well established at birth, whereas GFP-positive fibers in the superficial white matter, corresponding to U-fibers, are largely absent at P0 and progressively increase postnatally. Quantification of the GFP signal ratio, defined as the U-fiber region relative to deep white matter (Fig. 5B), confirms a significant increase from early to later stages (p < 0.01), indicating that U-fibers represent a late-forming fiber population.

**Fig. 5.**
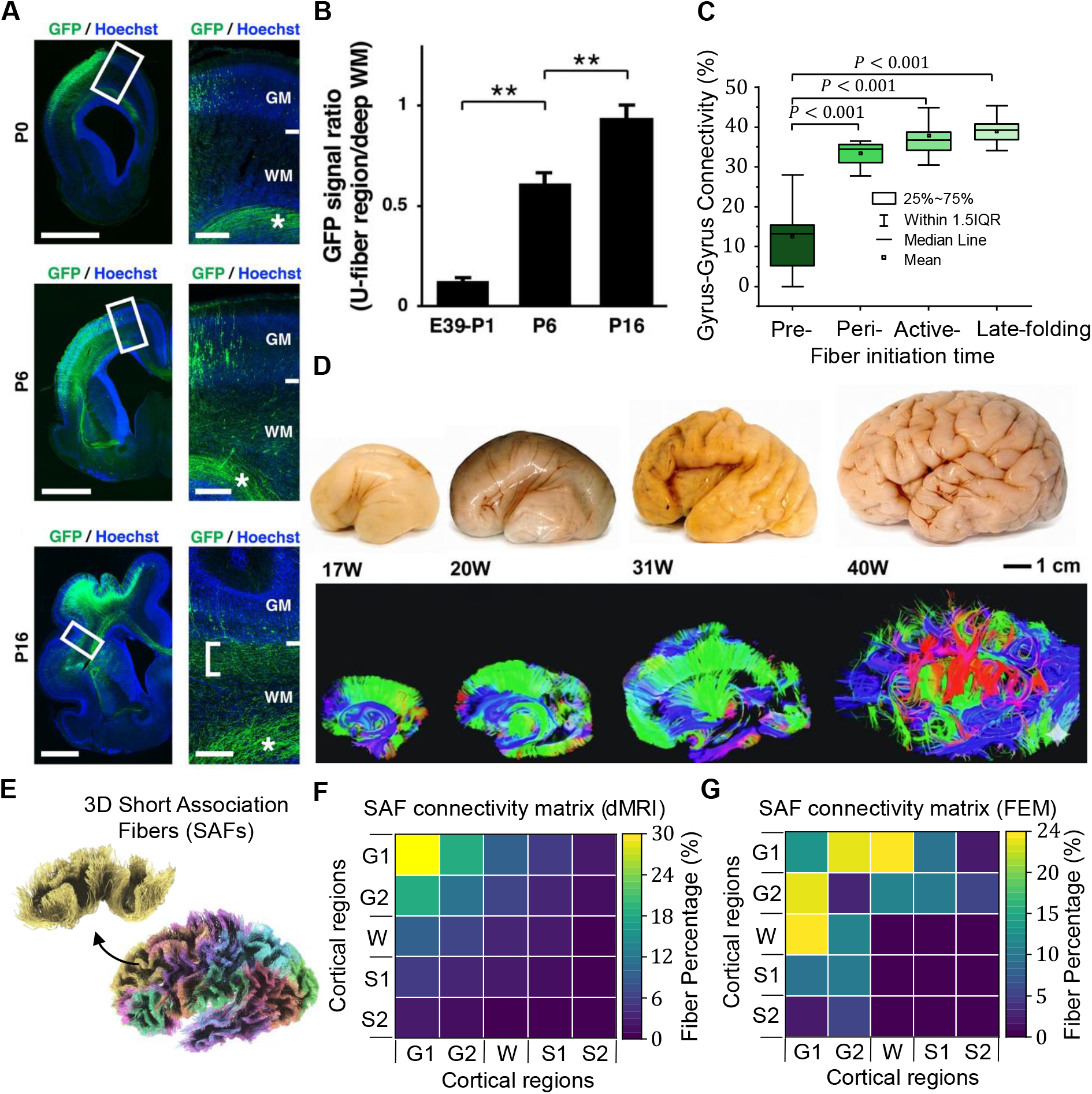
Multi-scale evidence supports a physical linkage between short association fiber (SAF) patterning and cortical folding. (**A–B**) Experimental observations of U-fiber development in ferret brains, reproduced from Yoshino et al.(*16*), “The origin and development of subcortical U-fibers in gyrencephalic ferrets,” *Mol Brain,* 2020, 13, 37, licensed under CC BY 4.0 (https://creativecommons.org/licenses/by/4.0/). GFP was introduced into cortical neurons via in utero electroporation at embryonic day (E31), and coronal sections were imaged at postnatal days P0, P6, and P16. (B) Quantification of the GFP signal ratio (U-fiber region/deep white matter) shows a significant increase from P1 to P16 (p < 0.01), indicating that U-fibers are a late-forming fiber population. Scale bars =2 mm (left) and 300 μm (right). (**C**) Model predictions of gyrus–gyrus connectivity across developmental stages (Pre, Peri, Active, and Late folding) show a marked increase after the onset of folding (post-bifurcation). (**D**) Human fetal brain tractography across gestational weeks (17–40 GW), illustrating the transition from dominant early radial pathways (17–20 GW) to progressively emerging corticocortical connections, including short-range association pathways and U-fibers (31–40 GW). Adapted from Takahashi et al.(*61*) “Emerging Cerebral Connectivity in the Human Fetal Brain: An MR Tractography Study,” *Cerebral Cortex*, 2012, 22(2): 455–464, by permission of Oxford University Press. Scale bar=1 cm. (**E**) Three-dimensional visualization of SAF streamline segments, highlighting distinct orientation groups and spatial organization within the superficial white matter. (**F–G**) Comparison of SAF connectivity matrices derived from dMRI and from the computational models. The data were extracted for the active-folding regime, and the parameters used in this figure are: *μ*_f_/*μ*_s_ = 2, *μ*_c_/*μ*_s_ = 2, *a* = 0.015 mm Pa^−1^ d^−1^, *G*^axn^ = 0.6 mm d^−1^, where *μ*_f_, *μ*_s_, and *μ*_c_ are the shear moduli of the fiber, subplate, and cortex, respectively, *a* is the stress-dependent elongation rate, and *G*^axn^ is the axon growth rate.

We next examined whether this delayed emergence is consistent with predictions from our mechanical model. Model derived gyrus-gyrus connectivity shown in Fig. 5C increases markedly after the onset of cortical folding, following bifurcation. Connectivity remains low during pre-folding stages and increases rapidly during active and late folding phases. This temporal trend closely mirrors the experimental observations in Fig. 5B, suggesting that short-range connections predominantly form after the establishment of folding patterns. The agreement between experiment and model supports the hypothesis that U-fiber-like connectivity is not pre-specified but emerges dynamically during cortical morphogenesis.

To place these findings in a human developmental context, we analyzed fetal brain tractography data (Fig. 5D), adapted from Takahashi et al.(*61*). Early stages at 17 to 20 GW are dominated by radial pathways, reflecting immature cortical organization. As development progresses to 31 to 40 GW, radial dominance decreases and corticocortical pathways, including short-range association fibers and U-fibers, become increasingly prominent, accompanied by more complex and crossing fiber architectures. This transition is consistent with the late-stage emergence of local connectivity observed in both experimental data and model predictions.

The structural organization of SAFs in a mature brain is further illustrated in Fig. 5E, where dMRI-derived streamline segments are visualized as color-coded sticks based on K-means clustering. This representation highlights distinct orientation groups and spatial organization within the superficial white matter, reflecting the organized yet heterogeneous arrangement of SAF pathways. To quantitatively assess connectivity patterns, we compared short-range connectivity matrices derived from dMRI and from our computational model (Fig. 5F-G). Both approaches exhibit enhanced local gyrus–gyrus connectivity, characterized by stronger near-diagonal elements, indicating preferential interactions between neighboring cortical regions.

## Discussion

This study introduced and evaluated a new mechanical theory for the formation, patterning, and establishment of the brain’s short association fibers (SAFs). While the structural and functional importance of SAFs are well-recognized, the mechanisms underlying their formation remain largely unexplored, particularly in terms of how biomechanical forces may shape their development during cortical folding and maturation. The results of this study provide a plausible mechanism for the formation and the characteristic U-shaped path of SAFs. In this context, the study’s findings are discussed below to illuminate their connections to existing literature and to highlight the insights they provide into the physical relationship between cortical folding and connectivity development.

### A new perspective on axonal tension theory

Recent advances in experimental and computational studies highlight the role of mechanical forces, alongside genetic and neural activity, in cortical development(*62–64*). Yet, the interplay between brain folding and connectivity development remains under debate(*65–67*). Current evidence suggests that integrating differential tangential growth and axonal tension offers a more comprehensive explanation for folding patterns and underlying connectivity observed in brain imaging studies(*22*, *23*, *29*, *42*, *68*). The traditional axonal tension theory(*34*) has evolved into a tension-induced subcortical growth model, proposing that subcortical axons respond to mechanical forces rather than initiate folding(*23*, *38*, *69*). Continuum-based models suggest folding stresses influence U-fiber patterns beneath sulci(*22*), while other models with explicit (non-growing) fibers show mechanical deformation alters fiber density across gyri and sulci(*42–44*). However, such models fall short in capturing dynamic connectivity development during brain folding(*14*, *70*).

A combined approach is essential to capture the response of individual, growing axonal fibers in response to mechanical stress fields induced by cortical folding. Our recent study(*45*) introduced a “stress-dependent axon reorientation” mechanism, simulating axons growing from deep regions into the folding cortex. Here, we extend this to axons emerging from the cortex to model cortico-cortical connectivity. The model enables fibers to exhibit stochastic, tip-driven, and stress-induced growth, either independently or concurrently, capturing diverse biological behaviors. As shown in Figs. 2, 3, S3, and S4, fibers dynamically reorient along tensile stress fields, aligning preferentially toward high-tension regions. Our simulations show cortical efferents navigating tangentially along sulci and radially near gyri. These stiff yet dynamic fibers may contribute to fold placement by forming a heterogeneous, evolving substrate for cortex folding(*29*, *42*). In the present model, intrinsic axonal growth is included, which may implicitly reflect underlying biochemical processes; however, reorientation is driven solely by stress-dependent mechanisms, and biochemical guidance(*71*) cues are not explicitly incorporated. It has been shown that in lissencephalic brains, such as those of rats and mice, where cortical folding is absent, SAFs can form through biochemical guidance mechanisms alone, although prominent U-shaped fiber systems are not developed(*72*). In contrast, as suggested by our model, cortical folding in gyrencephalic brains introduces heterogeneous mechanical stress fields that may provide an additional layer of guidance and contribute to the emergence of organized U-fiber structures.

We propose this as a potential mechanism for SAF formation, explaining why fibers emerging from the cortex adopt U-shaped paths and veer toward adjacent gyri rather than deeper brain structures in gyrencephalic brains (Fig. 2). Axonal stretching generates tension, aligning with experiments showing axons remain under tension during and after growth(*48*, *73*). These results correspond with stress patterns in the developing distinct brain(*38*) and DTI data from rhesus macaques(*22*). Our findings challenge both the traditional axonal tension theory(*34*) and recent axonal pushing theories(*74*, *75*), but support studies emphasizing ECM-generated mechanical cues in directing axonal growth(*76*). Additionally, our results closely mirror previous continuum-based models of axon growth and remodeling(*22*), which demonstrated that tension-induced remodeling within the developing subplate is sufficient to generate tissue-level anisotropic fiber orientations consistent with U-fibers. This mechanobiological perspective highlights the critical role of stress fields in shaping SAF connectivity during cortical development.

### Role of folding-connectivity association timing in SAF formation

During the second trimester, short-range functional connectivity strengthens with gestational age(*77*). SAFs begin forming mid to late gestation alongside cortical folding and mature postnatally(*51*, *61*, *78*). This timeline aligns with their role in functional network integration, essential for cognitive and sensory functions(*15*). We investigated how this timing affects SAF connectivity and found that fibers growing before folding mostly become projection fibers, while those initiating after folding are shaped by subplate stress fields and form U-fibers (Fig. 3). This may explain why, in gyrencephalic ferrets, U-fibers develop later than deeper white matter axons, as they likely respond to the stress field generated by cortical folding (Fig. 5A-B)(*16*). This observation is consistent with the emergence of SAFs following the initial growth of long-range projection fibers in the brain(*79*, *80*). The proposed stress-dependent axon reorientation mechanism is further supported by recent in utero imaging studies of SAF development(*52*). These studies show that SAFs do not initially exhibit U-shaped configurations, but instead emerge as relatively flat and loosely organized superficial pathways during early gestation, followed by rapid reorganization during the active folding phase, and eventual formation of coherent U-shaped bundles in later gestational stages. This progressive transition closely mirrors the behavior observed in our simulations, where early-emerging fibers are weakly influenced by the mechanical environment, while fibers emerging during and after folding undergo pronounced reorientation driven by evolving stress fields. The consistency between model predictions and the temporal progression of SAF development provides indirect but compelling support for the hypothesis that mechanical forces associated with cortical folding contribute to shaping short-range connectivity patterns, including the formation of U-fibers.

Studies in the gyrencephalic ferret brain have demonstrated the presence of stress fields consistent with our model during active cortical folding(*38*). The same study also examined stresses in the adult ferret, finding that similar stresses are maintained into maturity, albeit with reduced magnitude. Similarly, in Yucatan minipigs, mature white matter displayed anisotropic tensile stresses aligned with axon orientation(*53*), supporting our prediction of U-shaped trajectories for fibers developing during or after folding. Our simulation results suggest that the association timing between efferent fiber emergence and onset of folding can significantly influence connectivity development. In the extreme case of lissencephaly, absent folding-generated forces may impair SAF formation, contributing to dysconnectivity(*81*, *82*). More subtle alterations and delays of cortical folding have also been observed in brains of preterm-born individuals(*83*, *84*). A recent study found reduced frontal but increased occipital short-range connectivity in preterm infants(*85*). In primates, the establishment of short-range connectivity, including U-fibers, is generally considered more favorable than long-range radial connections due to wiring cost and efficiency constraints, and may support local cortical integration associated with higher-order cognitive functions(*86*, *87*). Higher local gyrification has also been linked to higher IQ and lower psychopathology in both preterm and control adults, though spatial patterns differ(*88*).

### Key biomechanical factors governing SAF formation and patterning

The present study also explored the theoretical effects of biomechanical factors on the development and organization of SAFs.

Simulations showed that faster axonal growth, or alternatively slower cortical growth, promotes more descending projection fibers and fewer tangentially oriented fibers (SAFs), especially beneath sulci (Fig. S3A-C). This matches observations in lissencephaly, where SAFs are reduced but projection fibers are preserved(*89*). Prior studies have shown that subplate-cortex growth dynamics affect folding: a faster subplate response reduces folding (lissencephaly/pachygyria), while a slower response leads to more small folds (polymicrogyria)(*23*). Both conditions are linked to reduced SAFs(*81*, *89*). In more subtle cases, such as preterm birth(*88*), reduced cortical expansion during folding aligns with our simulation predictions of reduced gyrus-gyrus connections (Fig. 4A), as seen in functional studies(*90*).

Our model also highlights the importance of cortex-to-subplate stiffness ratio in regulating folding timing and connectivity. Between 22–38 GW, brain tissue mechanics change substantially. Initially softer, white matter stiffens with gestation and after term due to myelination(*91*) and astrocytic network formation(*92*). Although gray matter is typically stiffer, the contrast is usually less than one order of magnitude(*59*, *93*, *94*). Measuring brain stiffness remains difficult due to its softness, structural complexity, and time-dependent behavior, including stress-dependent growth that reduces stiffness over time(*68*). Our simulations show that reduced stiffness contrast lowers SAF density (Figs. 4B and S4), supporting theories that such contrasts influence connectivity. This finding aligns with discussions(*92*) suggesting that early myelination, which begins during neurogenesis in rodents(*95*), increases white matter stiffness and may help explain why rodent brains exhibit less folding than those of larger mammals.

Finally, we investigated the impact of fiber stiffness on cortical folding and SAF formation. Most studies suggest that axons are stiffer than their surrounding substance as ECM(*96–99*). However, the stiffness difference spans an order of magnitude and is highly dependent on the testing methods or modeling assumptions employed. Our results suggest that variations in white matter stiffness, arising from the progressive growth of individual stiff fibers, can impact cortical fold placement. In line with past simulations of passive (non-growing fibers), gyri are more likely to emerge in regions with a higher density of stiff fibers(*74*, *75*, *100*), while sulci tend to form in areas with lower fiber density (Fig. S5). As shown in Fig. 4C, when the fiber-to-subplate stiffness ratio is 1, resulting in an isotropic substrate for cortical folding, fibers have no influence on the morphology of the folds. In contrast, as the stiffness ratio increases, stiffness variations arise within the subcortical domain, leading to alterations in fold morphology(*42*). This observation further supports the notion that brain connectivity may regulate and shape cortical folding patterns(*29*, *42*, *69*, *101*).

### Why are gyrus-gyrus connections more prevalent in SAFs?

Both tractography and histological studies generally indicate that SAFs predominantly connect neighboring gyri, whereas sulcal regions exhibit comparatively lower fiber density(*1*, *102*). Our imaging and simulation results support this, demonstrating more frequent gyrus-to-gyrus connections than other types (Figs. 3E, S3B, S4E, and 5G).

Our models indicate that the predominance of gyrus-to-gyrus connections arises from the orientation of MTPS in the subplate, shaped by cortical folding. Beneath gyri, MTPS is radially directed, promoting radial fiber growth through the gyral crown. In contrast, tangential MTPS beneath sulci steers fibers away from the sulcal cortex and toward adjacent gyri, where they align radially under tensile stress. This stress-guided mechanism favors the formation of connections between adjacent gyri rather than between gyri and sulci.

The multi-scale comparisons presented in Fig. 5 provide converging evidence that short-range connectivity, including U-fibers, emerges predominantly during the later stages of cortical development and is tightly coupled to the progression of folding. Experimental observations at the axonal level in gyrencephalic ferrets demonstrate that U-fibers are largely absent at early stages and increase significantly postnatally, indicating that they are a late-forming fiber population(*16*). Consistent with this, human fetal tractography studies reveal a transition from early radial organization to increasingly complex corticocortical architectures, with the progressive emergence of short-range association pathways and U-fibers during late gestation(*61*). At the network level, connectomic analyses further show an increase in local connectivity and functional segregation during this developmental window(*103*). Importantly, our model reproduces this delayed emergence, predicting a marked increase in gyrus–gyrus connectivity only after the onset of folding. The agreement across experimental, imaging, and computational observations suggests that the formation of SAF is not solely dictated by early genetic patterning, but is strongly influenced by the evolving physical environment of the cortex. In particular, the temporal coincidence between folding-induced stress fields and the increase in local connectivity supports a mechanistic role for stress-dependent axonal reorientation in shaping short-range pathways.

While the model captures the overall connectivity trends observed in tractography, discrepancies in connectivity patterns may arise (Fig. 5F-G). These differences likely reflect both model simplifications and inherent limitations of diffusion MRI, particularly the well-known gyral bias(*102*, *104*). Tractography is known to exhibit systematic inaccuracies in regions of complex geometry, especially within superficial white matter, where streamlines tend to terminate preferentially at gyral crowns and are underrepresented in sulcal regions. Accordingly, diffusion MRI is used here for qualitative comparison rather than as ground truth. The results are instead interpreted alongside histological observations from ferret and human brain development, which qualitatively support the predicted evolution of U fibers. At the same time, these limitations highlight the need for future studies that integrate high resolution histology and controlled experimental systems to further validate the proposed model.

### Limitations and future work

Modeling complex systems like the brain requires assumptions and simplifications, which naturally introduce limitations that point to future research directions.

First, mechanical parameters used in this study (e.g., axonal growth rate, cortex and fiber stiffnesses) were drawn from various sources, many of which may not reflect *in vivo* brain conditions during folding. For this reason, the analysis emphasized parametric and sensitivity studies rather than reliance on specific parameter values. A further limitation is the lack of quantitative *in vivo* or *ex vivo* measurements of axonal reorientation dynamics in response to tissue stress fields, which prevents direct calibration of the reorientation timescale and related model parameters. Together, these limitations highlight the need for targeted experimental studies on developing brain tissue, particularly those that directly probe axon reorientation under controlled mechanical environments, to refine model parameters and validate the proposed mechanisms.

Second, fiber tractography, while useful for mapping white matter, has known biases, especially in detecting radial fibers in gyri and resolving complex folding patterns(*102*). Recent studies integrating tractography with histological data support its utility while underscoring the need for multimodal approaches(*105–107*). However, additional histological studies focused specifically on SAFs are needed to accurately characterize their organization and connectivity, and to provide ground truth for validating fiber tractography and simulation models. A related limitation is that no new SAF-specific histological data were generated in the present study. The published ferret and human developmental datasets used for comparison were not acquired under experimental conditions directly matched to those represented in the model. Therefore, the agreement between these observations and the model predictions should be interpreted as corroborative rather than confirmatory evidence.

Finally, the model in this study was implemented in a 2D framework with simplified geometry, whereas brain growth and folding are inherently 3D processes. We deliberately adopted a 2D setup to enable hypothesis-driven analysis, reduce the number of parameters, and isolate the fundamental mechanisms governing stress-dependent axonal reorientation. In this context, the proposed mechanism depends on the local principal tensile stress direction, which is a tensorial quantity defined at each material point and exists independently of dimensionality. The essential requirement is the presence of anisotropic stress fields generated by differential growth, which are captured in both 2D and 3D formulations. While 3D effects may influence the detailed organization of fiber pathways, the core mechanism remains governed by local stress fields. However, it should be noted that the quantitative connectivity ratios reported in the 2D model, such as the percentages of gyrus-gyrus and gyrus-sulcus connections, should be interpreted as relative trends across developmental timing and biomechanical conditions rather than as absolute predictions for the three-dimensional brain. In a fully 3D cortical geometry, out-of-plane trajectories, gyral branching, regional curvature, and spatially heterogeneous folding patterns may alter the absolute values of these connectivity measures. Extending the present model to fully resolved 3D geometries remains an important direction for future work.

## Methods

### Constitutive framework

To model the growth and folding of the human brain, we adopt the differential tangential growth (DTG) theory as our foundational framework. Within this context, we highlight the dynamic interaction of efferent fibers with mechanical cues and cortical folding, as they grow and adapt as key structural components. We used the multiplicative decomposition of the deformation gradient(*108*) to separate the elastic and growth tensors. Based on this formulation, we developed constitutive equations describing tangential growth in the cortex and isotropic growth in the subcortex, both modeled using hyperelastic material behavior.

#### Multiplicative decomposition of the deformation gradient

As in continuum mechanics, we use the multiplicative decomposition of the total deformation gradient tensor **F**, where **F** accounts for both elastic (**F**^e^) and growth-induced deformation (**F**^g^). This approach enables us to separately characterize the reversible (elastic) and irreversible (growth-related) components of deformation:

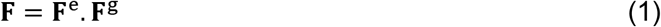

#### Cortical growth model

During the initial stages of cortical development, thickness variations are prominent(*109*); however, cortical folding is largely driven by changes in the surface area at later stages(*110*). This indicates that cortical growth predominantly occurs tangentially to the cortical surface, as noted in various studies(*111*). Thus, cortical growth is primarily modeled as in-plane area expansion. The growth tensor for the cortical layer is represented as:

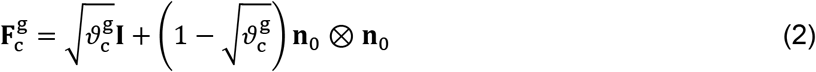

where 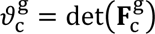 represents the area growth multiplier, **n**_0_ is the unit normal vector to the cortical surface, and **I** is the identity tensor. The corresponding growth rate follows a linear kinetic model:

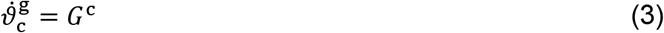

where *G*^c^ denotes the cortical growth rate. Then the elastic deformation gradient can be calculated by 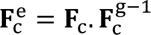.

#### Subcortex constitutive model

In contrast to the previous studies to model the white matter as a stretch-induced axonal growth media(*29*, *40*, *68*), we separated the subcortical region (later white matter) into two distinct parts as subplate and axon (fiber), as shown in Fig. 6A. We considered the subplate to represent the extracellular matrix (ECM) of the white matter, primarily composed of glial cells such as astrocytes and oligodendrocytes, along with structural proteins including glycoproteins and proteoglycans. The separation between the subplate and axonal parts provides an opportunity to explicitly model both the growth of the subplate and the reorientation of fibers. Observations indicate that white matter volume increases during development, so we assume isotropic growth of the subplate:

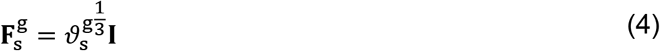

where **I** denotes the second order unit tensor and 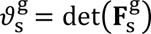 is the growth parameter represents the increase in the volume of the subplate. We define the growth rate as:

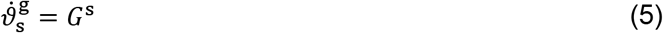

**Fig. 6.**
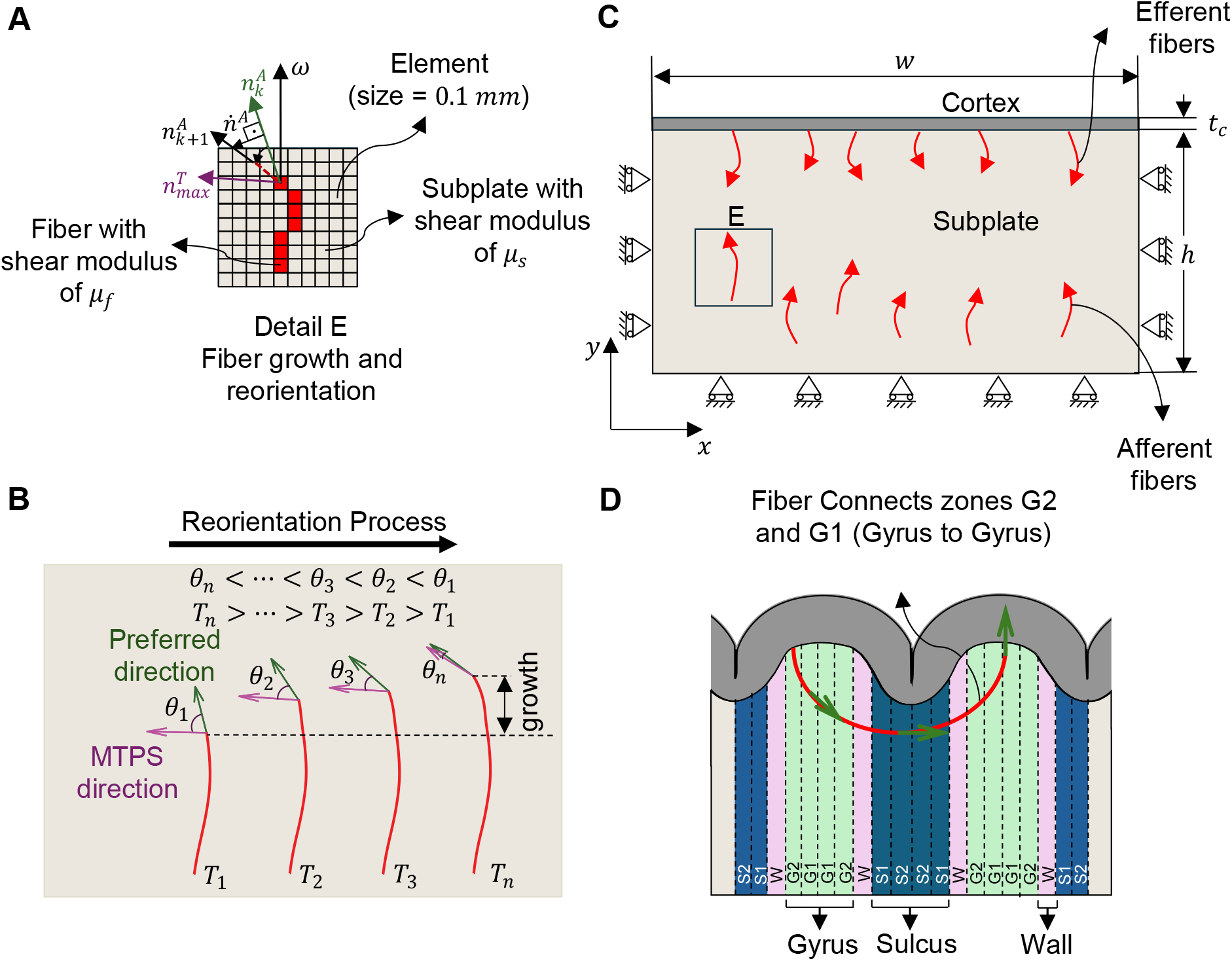
Schematic representation of the proposed model illustrating fiber growth and reorientation within the stress field of the subplate. (**A**) Illustration of fiber growth and navigation through subplate elements, with transformed material properties represented by red elements. (**B**) The fiber reorientation process over simulation time, showing a gradual alignment of the preferred direction toward the maximum tensile principal stress (MTPS) direction as time progresses. (**C**) Geometry and parameters of the finite element model: The model dimensions are defined as follows: width (*w*) of 60 mm, subplate thickness (ℎ) of 30 mm, and cortical thickness (*t_c_*) of 1.5 mm. (**D**) Division of the cortical plate (gyral span) into five equal-area bins, spanning from sulci to gyri, to identify connectivity streamlines linking these regions. The gyral span is the geodesic trajectory along the gray–white matter interface connecting the gyral crown to the sulcal fundus.

We model the subplate using neo-Hookean hyperelastic material, expressed as:

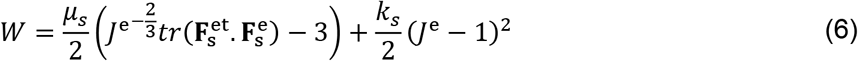

here, *μ_s_* and *k_s_* are shear and bulk moduli, and *J*^e^ is the elastic volume ratio. From this, the Cauchy stress tensor **T** is derived as:

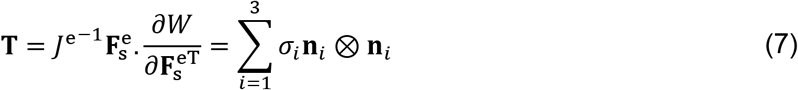

This allows to track principal stresses and directions, with directions **n***_i_* sorted by magnitude of principal stresses *σ*_1_ > *σ*_2_ > *σ*_3_. We also considered a neo-Hookean material for fibers with different shear modulus of *μ_f_* (Fig. 6A).

#### Stress-dependent axon reorientation

We updated our previously proposed method(*45*) for axon reorientation under mechanical loading to incorporate the growth of fibers. Fibers dynamically reorient along the principal stress directions, responding primarily to positive principal stress(*112*). The preferred direction **n**^A^ of each fiber aligns with the maximum tensile principal stress (MTPS) direction 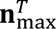 (Fig. 6B):

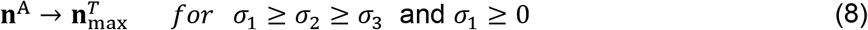

The rotational vector **ω**, given by:

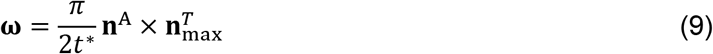

where *t*^∗^ is a relaxation parameter, controls the reorientation rate. This vector is decomposed as **ω** = *ω***n***^ω^* with **n***^ω^* as the unit normal vector of **ω**. The variation of the preferred direction over time can be depicted as a rotation about axis *ω* as shown in Fig. 6A. Therefore, the evolution of the preferred direction over time is expressed as follows:

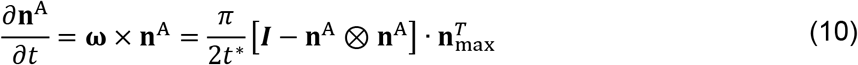

The Preferred direction, **n**^A^, changes over time at a rate, 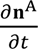, influenced by 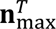. This change, denoted as **n***_A_* in Fig. 6A, which is perpendicular to **n**^A^. This rate is scaled by 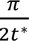, resulting in reorientation that is proportional to the sinus of the angle between **n**^A^ and 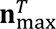. We used a discrete model to solve Eq. (10) and capture the evolution of fiber directions. The details of this discrete model are provided in the Methods section of the Supplementary Materials and in our previous publication(*45*).

#### Tip and stress-induced axon growth

Our model incorporates two key mechanisms of axon elongation: tip-driven growth and stress-induced stretching. Tip-driven, or intrinsic, growth occurs at the axon’s growth cone and depends on biochemical cues and nutrient availability, allowing elongation to proceed independently of mechanical forces if those conditions are met(*113*). In contrast, stress-induced growth, also known as stretch-dependent or towed growth, results from mechanical tension along the axon shaft due to deformations in the extracellular matrix(*39*). To capture both mechanisms, the model integrates a baseline fixed growth rate, *G*^axn^, with a stress-dependent elongation term, allowing axons to grow directionally in response to both internal and external cues. The total growth rate is defined as:

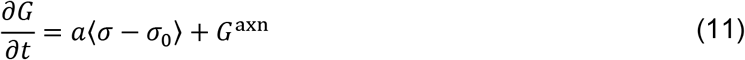

where *σ* is the maximum tensile principal stress along the axon’s orientation, *σ*_0_ is the homeostatic stress level (set to zero in our model), and *a* is the sensitivity coefficient for stress-induced growth. The Macauley bracket ⟨⋅⟩ ensures that only tensile stresses exceeding the threshold *σ*_0_ contribute to growth, while compressive stresses do not influence the growth rate. When *σ* ≤ *σ*_0_, the growth rate reduces to the baseline term *G*^axn^, and axonal growth proceeds in a stochastic manner. This formulation reflects experimental findings that embryonic axons, such as those in the forebrain, can undergo significant elongation in response to even minimal tension(*114*). Details of the dynamic framework and the algorithm update scheme for stress-dependent axon reorientation are provided in the Methods section of the Supplementary Materials.

### Finite element model

The finite element (FE) model simulates brain development as a bilayer system composed of a tangentially expanding cortical layer and an underlying subcortical layer made of subplate and growing fibers (Fig. 6C). Implemented in Abaqus/Explicit using a custom material subroutine, the model includes realistic dimensions, meshing, and boundary conditions that replicate cortical folding driven by differential growth. While the focus of the study is on efferent fibers, afferent fibers were also modeled in the FE models to account for their potential influence on the patterning of efferent fibers. However, the results indicate that the inclusion of afferent fibers does not affect the behavior or developmental characteristics of SAFs, as shown in Fig. S2. Therefore, fiber elements represent either afferent or efferent axons with distinct origins and growth directions. The model tracks fiber growth dynamically by updating their orientation toward the direction of maximum tensile principal stress (MTPS) during each time step. The algorithm reclassifies subplate elements into fibers as growth progresses, and uses a relaxation parameter to ensure smooth reorientation, aligning with biological observations.

Fiber elongation is governed by a fixed axonal growth rate plus a stress-dependent component, with axons growing faster and reorienting more precisely under mechanical stress. A nonlinear hyperelastic material model captures the elastic and stretch-responsive behavior of axons during deformation. Baseline shear modulus ratios for cortex-to-subplate and axon-to-subplate are set at 2, reflecting conditions in the developing brain with low myelination. Literature consistently indicates that axonal fibers are stiffer than the surrounding ECM, with reported stiffness ratios varying depending on measurement conditions or micromechanical modeling approache(*59*, *115*). Considering that axons in the fetal brain are not yet fully myelinated, we adopted a conservative shear modulus ratio of axonal fibers to ECM of 2 in the present study. Our prior sensitivity analyses further indicate that the model predictions remain insensitive to the exact value of this ratio, provided that axonal fibers are relatively stiffer than the surrounding ECM(*45*). The model assumes varying axonal growth rates and provides a parametric platform for exploring how mechanical properties and growth dynamics influence folding patterns and white matter organization. A complete list of model parameters is provided in Table 1. More details of the FE model, including the rationale behind the selection of model parameters, are provided in the Methods section of the Supplementary Materials.

**Table 1.** Parameters and their values used in the simulations.

| Parameter | Cortical thickness | Simulation duration | Cortical growth rate | Axon growth rate | Stiffness ratio of cortex to subplate | Stiffness ratio of fibers to subplate | stress-dependent axon elongation rate | Relaxation parameter |
| --- | --- | --- | --- | --- | --- | --- | --- | --- |
| | (mm) | days (d) | $G^c$ ($\text{d}^{-1}$) | ($\text{mm d}^{-1}$) | ($\mu_c/\mu_s$) | ($\mu_f/\mu_s$) | $a$ ($\text{mm Pa}^{-1} \text{d}^{-1}$) | $t^*$ |
| Values | 1.5 | 50 | 0.02 | 0.6, 0.8, 1.0, 1.2 | 1, 2, 3, 4 | 1, 2, 3, 4 | 0.015 | 200 |

All simulations were performed over *t* ∈ [0,50] days, corresponding to the normalized model interval *T* ∈ [0,1] through *T* = *G^c^t*. This parameterization, retained from our previously established modeling frameworks(*22*, *45*), provides a consistent computational timescale for controlled comparisons across the different timing regimes. The 50-day interval was not calibrated to represent the complete 22–38 GW period of human fetal development.

The relaxation parameter *t*^∗^ and *a* were determined from the sensitivity analysis presented in Fig. S7. All quantitative analyses are based on simulations with 150 fibers per model. A convergence analysis shows that increasing the number of fibers beyond approximately 150 has negligible impact on key metrics, including connectivity patterns, indicating statistical stabilization (Fig. S8A). For visualization, a reduced set of 100 fibers is shown to improve clarity in illustrating fiber trajectories and their interaction with the evolving cortical geometry. To minimize dependence on fiber initiation positions, a sensitivity analysis was performed, demonstrating that reliable statistical results are obtained by averaging over 10 independent trials (Fig. S8B). Accordingly, all simulations were conducted using 10 trials per parameter set, and results are reported as mean values with corresponding standard deviations.

#### Distribution of connections along the gyral span

We define the gyral span as the geodesic trajectory along the gray–white matter interface connecting the gyral crown to the sulcal fundus. The distribution of SAFs within the gyral span, in relation to gyri, sulci, and cortical walls, was analyzed by dividing it into five equal-area zones extending from the sulcal fundus to the gyral crown, following the methodology outlined by Cottaar et al.(*116*). This segmentation, illustrated in Fig. 6D, classified the middle zone as the cortical wall, while the remaining zones were designated as either gyrus or sulcus regions. Connectivity matrices were then constructed based on the fibers counts, providing a detailed representation of fiber connectivity distribution. We applied the same methodology for dMRI data to partition the cortical surface into five equidistant regions and constructed a 5×5 connectivity matrix. Furthermore, a group-level average connectivity matrix was computed to capture the overall connectivity pattern.

### Brain imaging data and processing

The brain imaging data and processing are described in detail in the Methods section of the Supplementary Materials. Here, we provide a brief overview of the key steps relevant to SAF extraction. In this study, we used grayordinate T1/2-weighted (T1/2w) MRI data and diffusion MRI (dMRI) data from the Human Connectome Project (HCP) 3T dataset, provided by the Washington University–Minnesota consortium(*117*). The participants were between 22 and 35 years of age, with an approximately balanced gender distribution. For our analysis, we randomly selected a cohort of 14 subjects from the HCP 3T dataset.

#### Ethics statement

This study did not involve the collection of new human subjects data. All imaging data were obtained from the publicly available, de-identified Human Connectome Project (HCP) dataset(*117*). The original HCP study protocols were approved by the institutional review board of Washington University in St. Louis, and all participants provided informed consent. Therefore, no additional ethical approval was required for this study.

#### Diffusion MRI preprocessing and short association fibers extraction

The diffusion MRI preprocessing and SAF extraction pipeline (Fig. 7) involved established tools such as MRtrix3.0(*118*) and ANTs(*119*), incorporating denoising, motion correction, and tissue segmentation from high-resolution structural images. Diffusion-weighted data were processed to estimate multi-shell, multi-tissue response functions and reconstruct fiber orientation distributions (FODs) using constrained spherical deconvolution. Whole-brain probabilistic tractography was performed with anatomical constraints, and SAFs were isolated by filtering streamlines shorter than 40 mm that connect neocortical gray matter regions within the same hemisphere. Streamlines terminating in subcortical regions or crossing hemispheres were excluded. Finally, short-range connectivity is quantified by mapping SAF endpoints to cortical regions, yielding region-wise connectivity matrices that capture the percentage of local (gyrus–gyrus) interactions.

**Fig. 7.**
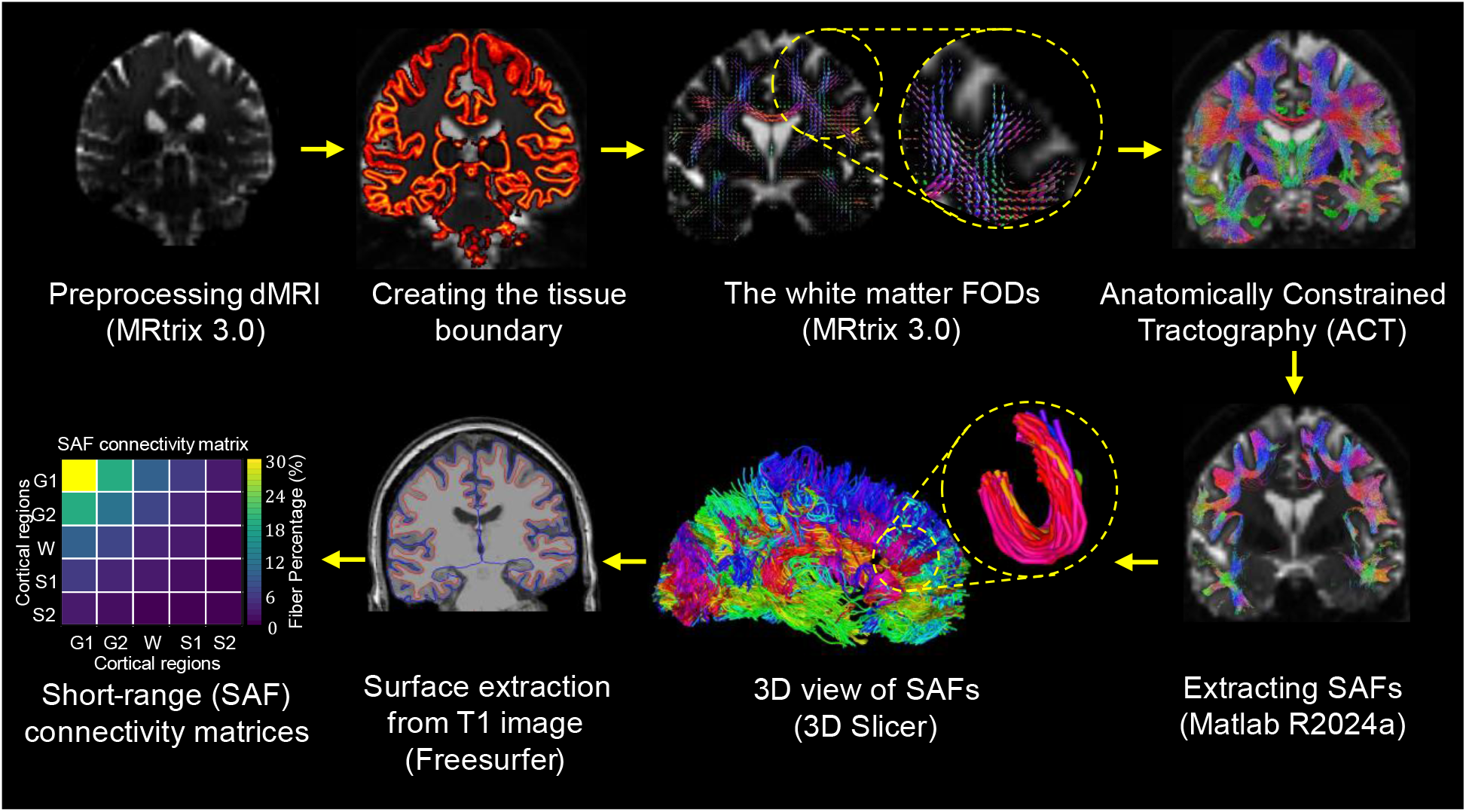
Overview of the processing pipeline for short association fibers (SAFs) extraction. The workflow begins with preprocessing of diffusion MRI data, followed by generation of tissue boundaries from structural T1-weighted images. White matter Fiber Orientation Distributions (FODs) are then estimated and used for Anatomically Constrained Tractography (ACT) to generate biologically plausible streamlines. A custom algorithm is applied to extract SAFs by selecting streamlines that connect adjacent cortical regions within the same hemisphere. A 3D view illustrates the resulting SAFs. Cortical surfaces are reconstructed from the T1-weighted image using FreeSurfer, and SAF endpoints are mapped to cortical regions to compute short-range connectivity matrices representing gyrus–gyrus interactions.

### Statistical analysis

We conducted a statistical analysis of our study findings using SPSS to evaluate the influence of biophysical properties of axons, the cortex, and the subplate on fiber organization. To compare multiple groups, we utilized Tukey’s Honestly Significant Difference (HSD) test with a significance threshold of 0.05 (95% confidence level). This method was particularly effective in identifying statistically significant differences between group pairs.

## Acknowledgments

The authors acknowledge that data were provided in part by the Human Connectome Project, WU-Minn Consortium (Principal Investigators: David Van Essen and Kamil Ugurbil; 1U54MH091657), funded by the 16 NIH Institutes and Centers that support the NIH Blueprint for Neuroscience Research, and by the McDonnell Center for Systems Neuroscience at Washington University. **Funding:** M. J. Razavi acknowledges support from the National Science Foundation (CMMI-2123061). K. E. Garcia acknowledges support from the National Institutes of Health (R01 NS133116). X. Wang acknowledges the support from National Science Foundation (IIS-2011369) and National Institutes of Health (1R01NS135574-01). **Author contributions:** A. S.: Writing – original draft, Investigation, Writing – review & editing, Methodology, Resources, Data curation, Validation, Supervision, Formal analysis, Software, Project administration, Visualization. Z. H.: Writing – original draft, Methodology, Formal analysis, Software. J. H.: Writing – original draft, Methodology, Formal analysis, Software. K. E. G.: Conceptualization, Writing – review & editing, Methodology. T. Z.: Investigation, Writing – review & editing, Methodology, Validation. L. P.: Conceptualization, Writing – review & editing, Resources. G. C.: Writing – original draft, Methodology, Formal analysis, Software. X. W.: Investigation, Writing – review & editing, Funding acquisition, Supervision. M. J. R.: Writing – original draft, Conceptualization, Investigation, Writing – review & editing, Methodology, Funding acquisition, Validation, Supervision, Project administration. All authors reviewed the manuscript. **Competing Interests:** The authors declare no competing interests. **Data, code, and materials availability:** All data and code needed to evaluate and reproduce the results in the paper are present in the paper, the Supplementary Materials, and/or deposited in Zenodo at https://doi.org/10.5281/zenodo.20099058. The Zenodo repository includes the model files, analysis codes, and processed MRI data used in this study. This study did not generate new materials.

## Supplementary Materials

This PDF file Includes: Supplementary Text Figs. S1 to S8

Table S1 References

## References

1. D. Shastin, S. Genc, G. D. Parker, K. Koller, C. M. W. Tax, J. Evans, K. Hamandi, W. P. Gray, D. K. Jones, M. Chamberland, Surface-based tracking for short association fibre tractography. NeuroImage 260, 119423 (2022).

2. G. Dannhoff, P. P. Poudel, C. Bhattarai, S. G. Kalthur, I. L. Maldonado, Depicting the anatomy of the gyral white matter: *ubi sumus? quo vadimus?* Brain Communications 5, fcad265 (2023).

3. O. R. Phillips, K. A. Clark, E. Luders, R. Azhir, S. H. Joshi, R. P. Woods, J. C. Mazziotta, A. W. Toga, K. L. Narr, Superficial White Matter: Effects of Age, Sex, and Hemisphere. Brain Connectivity 3, 146–159 (2013).

4. M. Wu, L. H. Lu, A. Lowes, S. Yang, A. M. Passarotti, X. J. Zhou, M. N. Pavuluri, Development of superficial white matter and its structural interplay with cortical gray matter in children and adolescents. Human Brain Mapping 35, 2806–2816 (2014).

5. M. Catani, F. Dell’Acqua, F. Vergani, F. Malik, H. Hodge, P. Roy, R. Valabregue, M. Thiebaut De Schotten, Short frontal lobe connections of the human brain. Cortex 48, 273–291 (2012).

6. M. Catani, N. Robertsson, A. Beyh, V. Huynh, F. De Santiago Requejo, H. Howells, R. L. C. Barrett, M. Aiello, C. Cavaliere, T. B. Dyrby, K. Krug, M. Ptito, H. D’Arceuil, S. J. Forkel, F. Dell’Acqua, Short parietal lobe connections of the human and monkey brain. Cortex 97, 339–357 (2017).

7. M. Wilkinson, R. Wang, A. Van Der Kouwe, E. Takahashi, White and gray matter fiber pathways in autism spectrum disorder revealed by ex vivo diffusion MR tractography. Brain and Behavior 6, e00483 (2016).

8. M. Liu, B. C. Bernhardt, S.-J. Hong, B. Caldairou, A. Bernasconi, N. Bernasconi, The superficial white matter in temporal lobe epilepsy: a key link between structural and functional network disruptions. Brain 139, 2431–2440 (2016).

9. R. O’Halloran, R. Feldman, L. Marcuse, M. Fields, B. Delman, S. Frangou, P. Balchandani, A method for u-fiber quantification from 7 T diffusion-weighted MRI data tested in patients with nonlesional focal epilepsy. NeuroReport 28, 457–461 (2017).

10. O. R. Phillips, S. H. Joshi, F. Piras, M. D. Orfei, M. Iorio, K. L. Narr, D. W. Shattuck, C. Caltagirone, G. Spalletta, M. Di Paola, The superficial white matter in Alzheimer’s disease. Human Brain Mapping 37, 1321–1334 (2016).

11. C. Carmeli, E. Fornari, M. Jalili, R. Meuli, M. G. Knyazeva, Structural covariance of superficial white matter in mild Alzheimer’s disease compared to normal aging. Brain and Behavior 4, 721–737 (2014).

12. H. C. Kinney, B. Ann Brody, A. S. Kloman, F. H. Gilles, Sequence of Central Nervous System Myelination in Human Infancy. II. Patterns of Myelination in Autopsied Infants: Journal of Neuropathology and Experimental Neurology 47, 217–234 (1988).

13. B. A. Brody, H. C. Kinney, A. S. Kloman, F. H. Gilles, Sequence of Central Nervous System Myelination in Human Infancy. I. An Autopsy Study of Myelination: Journal of Neuropathology and Experimental Neurology 46, 283–301 (1987).

14. I. Kostović, M. Radoš, M. Kostović-Srzentić, Ž. Krsnik, Fundamentals of the Development of Connectivity in the Human Fetal Brain in Late Gestation: From 24 Weeks Gestational Age to Term. Journal of Neuropathology & Experimental Neurology 80, 393–414 (2021).

15. M. Ouyang, H. Kang, J. A. Detre, T. P. L. Roberts, H. Huang, Short-range connections in the developmental connectome during typical and atypical brain maturation. Neuroscience & Biobehavioral Reviews 83, 109–122 (2017).

16. M. Yoshino, K. Saito, K. Kawasaki, T. Horiike, Y. Shinmyo, H. Kawasaki, The origin and development of subcortical U-fibers in gyrencephalic ferrets. Mol Brain 13, 37 (2020).

17. H. Huang, J. Zhang, S. Wakana, W. Zhang, T. Ren, L. J. Richards, P. Yarowsky, P. Donohue, E. Graham, P. C. M. Van Zijl, S. Mori, White and gray matter development in human fetal, newborn and pediatric brains. NeuroImage 33, 27–38 (2006).

18. J. Dubois, G. Dehaene-Lambertz, S. Kulikova, C. Poupon, P. S. Hüppi, L. Hertz-Pannier, The early development of brain white matter: A review of imaging studies in fetuses, newborns and infants. Neuroscience 276, 48–71 (2014).

19. G. Zheng, B. Fei, A. Ge, Y. Liu, Y. Liu, Z. Yang, Z. Chen, X. Wang, H. Wang, J. Ding, U-fiber analysis: a toolbox for automated quantification of U-fibers and white matter hyperintensities. Quant Imaging Med Surg 14, 662–683 (2024).

20. E. Kirilina, S. Helbling, M. Morawski, K. Pine, K. Reimann, S. Jankuhn, J. Dinse, A. Deistung, J. R. Reichenbach, R. Trampel, S. Geyer, L. Müller, N. Jakubowski, T. Arendt, P.-L. Bazin, N. Weiskopf, Superficial white matter imaging: Contrast mechanisms and whole-brain in vivo mapping. Sci. Adv. 6, eaaz9281 (2020).

21. K. E. Garcia, E. C. Robinson, D. Alexopoulos, D. L. Dierker, M. F. Glasser, T. S. Coalson, C. M. Ortinau, D. Rueckert, L. A. Taber, D. C. Van Essen, C. E. Rogers, C. D. Smyser, P. V. Bayly, Dynamic patterns of cortical expansion during folding of the preterm human brain. Proc Natl Acad Sci USA 115, 3156–3161 (2018).

22. K. E. Garcia, X. Wang, C. D. Kroenke, A model of tension-induced fiber growth predicts white matter organization during brain folding. Nat Commun 12, 6681 (2021).

23. P. V. Bayly, R. J. Okamoto, G. Xu, Y. Shi, L. A. Taber, A cortical folding model incorporating stress-dependent growth explains gyral wavelengths and stress patterns in the developing brain. Phys. Biol. 10, 016005 (2013).

24. P. V. Bayly, L. A. Taber, C. D. Kroenke, Mechanical forces in cerebral cortical folding: A review of measurements and models. Journal of the Mechanical Behavior of Biomedical Materials 29, 568–581 (2014).

25. S. Budday, P. Steinmann, E. Kuhl, The role of mechanics during brain development. Journal of the Mechanics and Physics of Solids 72, 75–92 (2014).

26. S. Budday, P. Steinmann, A. Goriely, E. Kuhl, Size and curvature regulate pattern selection in the mammalian brain. Extreme Mechanics Letters 4, 193–198 (2015).

27. T. Tallinen, J. Y. Chung, J. S. Biggins, L. Mahadevan, Gyrification from constrained cortical expansion. Proceedings of the National Academy of Sciences 111, 12667–12672 (2014).

28. T. Tallinen, J. Y. Chung, F. Rousseau, N. Girard, J. Lefèvre, L. Mahadevan, On the growth and form of cortical convolutions. Nature Phys 12, 588–593 (2016).

29. M. A. Holland, K. E. Miller, E. Kuhl, Emerging Brain Morphologies from Axonal Elongation. Ann Biomed Eng 43, 1640–1653 (2015).

30. S. Wang, N. Demirci, M. A. Holland, Numerical investigation of biomechanically coupled growth in cortical folding. Biomech Model Mechanobiol 20, 555–567 (2021).

31. M. J. Razavi, T. Zhang, T. Liu, X. Wang, Cortical Folding Pattern and its Consistency Induced by Biological Growth. Scientific Reports 5 (2015).

32. M. J. Razavi, T. Zhang, X. Li, T. Liu, X. Wang, Role of mechanical factors in cortical folding development. *Phys*. Rev. E 92, 032701 (2015).

33. D. P. Richman, R. M. Stewart, J. Hutchinson, V. S. Caviness, Mechanical Model of Brain Convolutional Development: Pathologic and experimental data suggest a model based on differential growth within the cerebral cortex. Science 189, 18–21 (1975).

34. D. C. V. Essen, A tension-based theory of morphogenesis and compact wiring in the central nervous system. Nature 385, 313–318 (1997).

35. D. C. Van Essen, A 2020 view of tension-based cortical morphogenesis. Proc. Natl. Acad. Sci. U.S.A. 117, 32868–32879 (2020).

36. A. Solhtalab, Y. Guo, A. Gholipour, W. Dai, M. J. Razavi, Mechanics of the Spatiotemporal Evolution of Sulcal Pits in the Folding Brain. Human Brain Mapping 46, e70332 (2025).

37. X. Wang, J. Hou, Z. Wu, K. Jiang, T. Wu, L. Zhang, D. Zhu, W. Gao, M. Razavi, T. Liu, E. Kuhl, G. Li, Biophysical modeling of anatomically realistic prenatal cortical folding development. Res Sq, rs.3.rs-8033969 (2026).

38. G. Xu, A. K. Knutsen, K. Dikranian, C. D. Kroenke, P. V. Bayly, L. A. Taber, Axons Pull on the Brain, But Tension Does Not Drive Cortical Folding. Journal of Biomechanical Engineering 132, 071013 (2010).

39. A. Goriely, S. Budday, E. Kuhl, “Neuromechanics” in Advances in Applied Mechanics (Elsevier, 2015; https://linkinghub.elsevier.com/retrieve/pii/S0065215615000034)vol. 48, pp. 79–139.

40. S. Budday, C. Raybaud, E. Kuhl, A mechanical model predicts morphological abnormalities in the developing human brain. Sci Rep 4, 5644 (2015).

41. A. H Foroughi, A. Solhtalab, G. Cao, T. Zhang, M. J. Razavi, A mechanical model for axon pathfinding in a folding brain. J. Phys. D: Appl. Phys. 58, 175401 (2025).

42. X. Wang, S. Wang, M. A. Holland, Axonal tension contributes to consistent fold placement. Soft Matter 20, 3053–3065 (2024).

43. P. Chavoshnejad, L. Vallejo, S. Zhang, Y. Guo, W. Dai, T. Zhang, M. J. Razavi, Mechanical hierarchy in the formation and modulation of cortical folding patterns. Sci Rep 13, 13177 (2023).

44. P. Chavoshnejad, X. Li, S. Zhang, W. Dai, L. Vasung, T. Liu, T. Zhang, X. Wang, M. J. Razavi, Role of axonal fibers in the cortical folding patterns: A tale of variability and regularity. Brain Multiphysics 2, 100029 (2021).

45. A. Solhtalab, A. H. Foroughi, L. Pierotich, M. J. Razavi, Stress landscape of folding brain serves as a map for axonal pathfinding. Nat Commun 16, 1187 (2025).

46. J. C. Silbereis, S. Pochareddy, Y. Zhu, M. Li, N. Sestan, The Cellular and Molecular Landscapes of the Developing Human Central Nervous System. Neuron 89, 248–268 (2016).

47. D. E. Koser, A. J. Thompson, S. K. Foster, A. Dwivedy, E. K. Pillai, G. K. Sheridan, H. Svoboda, M. Viana, L. D. F. Costa, J. Guck, C. E. Holt, K. Franze, Mechanosensing is critical for axon growth in the developing brain. Nat Neurosci 19, 1592–1598 (2016).

48. K. Franze, Integrating Chemistry and Mechanics: The Forces Driving Axon Growth. Annu. Rev. Cell Dev. Biol. 36, 61–83 (2020).

49. H. de Vareilles, D. Rivière, J. F. Mangin, J. Dubois, Development of cortical folds in the human brain: An attempt to review biological hypotheses, early neuroimaging investigations and functional correlates. Dev Cogn Neurosci 61, 101249 (2023).

50. G. M. Innocenti, D. J. Price, Exuberance in the development of cortical networks. Nat Rev Neurosci 6, 955–965 (2005).

51. I. Kostovic, P. Rakic, Developmental history of the transient subplate zone in the visual and somatosensory cortex of the macaque monkey and human brain. J of Comparative Neurology 297, 441–470 (1990).

52. B. Li, Ž. Krsnik, L. Pierotich, I. Kostović, S. K. Warfield, P. E. Grant, D. Karimi, Insights into Temporal and Spatial Dynamics of Short Association Fiber Formation in the Human Fetal Brain. Neuroscience [Preprint] (2025). 10.1101/2025.10.20.683525.

53. R. Balouchzadeh, C. D. Kroenke, K. E. Garcia, P. V. Bayly, Measurement of residual stress in the brain. [Preprint] (2025). 10.1101/2025.02.26.637925.

54. D. K. Shukla, B. Keehn, D. M. Smylie, R.-A. Müller, Microstructural abnormalities of short-distance white matter tracts in autism spectrum disorder. Neuropsychologia 49, 1378–1382 (2011).

55. S. Zhang, Y. Wang, F. Deng, S. Zhong, L. Chen, X. Luo, S. Qiu, P. Chen, G. Chen, H. Hu, S. Lai, H. Huang, Y. Jia, L. Huang, R. Huang, Disruption of superficial white matter in the emotion regulation network in bipolar disorder. NeuroImage: Clinical 20, 875–882 (2018).

56. J. Hou, Z. Wu, X. Chen, L. Wang, D. Zhu, T. Liu, G. Li, X. Wang, Role of data-driven regional growth model in shaping brain folding patterns. Soft Matter 21, 729–749 (2025).

57. T. Zhang, M. J. Razavi, X. Li, H. Chen, T. Liu, X. Wang, Mechanism of Consistent Gyrus Formation: an Experimental and Computational Study. Sci Rep 6, 37272 (2016).

58. F. Atashgar, M. Shafieian, N. Abolfathi, From structure to mechanics: exploring the role of axons and interconnections in anisotropic behavior of brain white matter. Biomech Model Mechanobiol, doi: 10.1007/s10237-025-01957-4 (2025).

59. S. Budday, G. Sommer, C. Birkl, C. Langkammer, J. Haybaeck, J. Kohnert, M. Bauer, F. Paulsen, P. Steinmann, E. Kuhl, G. A. Holzapfel, Mechanical characterization of human brain tissue. Acta Biomaterialia 48, 319–340 (2017).

60. J. Hou, K. Jiang, A. Ramanathan, A. S. Kumar, W. Zhang, L. Zhao, T. Wu, R. Pidaparti, D. Zhu, G. Li, K. Song, T. Liu, M. J. Razavi, E. Kuhl, X. Wang, Mechanical Characterization of Brain Tissue: Experimental Techniques, Human Testing Considerations, and Perspectives. arXiv arXiv:2504.12346 [Preprint] (2025). 10.48550/arXiv.2504.12346.

61. E. Takahashi, R. D. Folkerth, A. M. Galaburda, P. E. Grant, Emerging Cerebral Connectivity in the Human Fetal Brain: An MR Tractography Study. Cerebral Cortex 22, 455–464 (2012).

62. K. E. Garcia, C. D. Kroenke, P. V. Bayly, Mechanics of cortical folding: stress, growth and stability. Phil. Trans. R. Soc. B 373, 20170321 (2018).

63. D. C. Van Essen, Biomechanical models and mechanisms of cellular morphogenesis and cerebral cortical expansion and folding. Seminars in Cell & Developmental Biology 140, 90–104 (2023).

64. C. Llinares-Benadero, V. Borrell, Deconstructing cortical folding: genetic, cellular and mechanical determinants. Nat Rev Neurosci 20, 161–176 (2019).

65. J. I. Arellano, P. Rakic, “Old Models Know Wrinkles Best: A Critical Review on the Mechanisms of Cortical Gyrification” in Neocortical Neurogenesis in Development and Evolution, W. Huttner, Ed. (Wiley, ed. 1, 2023; https://onlinelibrary.wiley.com/doi/10.1002/9781119860914.ch23), pp. 499–525.

66. P. V. Bayly, Perspective: Challenges and opportunities in computational brain mechanics research: How can we use recent experimental data to improve models of brain mechanics? Brain Multiphysics 4, 100075 (2023).

67. S. Budday, Exploring human brain mechanics by combining experiments, modeling, and simulation. Brain Multiphysics 5, 100076 (2023).

68. R. Balouchzadeh, P. V. Bayly, K. E. Garcia, Effects of stress-dependent growth on evolution of sulcal direction and curvature in models of cortical folding. Brain Multiphysics 4, 100065 (2023).

69. K. E. Garcia, C. D. Kroenke, P. V. Bayly, Mechanical stress connects cortical folding to fiber organization in the developing brain. Trends in Neurosciences, S0166223625000761 (2025).

70. M. E. Thomason, Development of Brain Networks In Utero: Relevance for Common Neural Disorders. Biol Psychiatry 88, 40–50 (2020).

71. M. O’Donnell, R. K. Chance, G. J. Bashaw, Axon Growth and Guidance: Receptor Regulation and Signal Transduction. Annu. Rev. Neurosci. 32, 383–412 (2009).

72. Q. Wang, S.-L. Ding, Y. Li, J. Royall, D. Feng, P. Lesnar, N. Graddis, M. Naeemi, B. Facer, A. Ho, T. Dolbeare, B. Blanchard, N. Dee, W. Wakeman, K. E. Hirokawa, A. Szafer, S. M. Sunkin, S. W. Oh, A. Bernard, J. W. Phillips, M. Hawrylycz, C. Koch, H. Zeng, J. A. Harris, L. Ng, The Allen Mouse Brain Common Coordinate Framework: A 3D Reference Atlas. Cell 181, 936–953.e20 (2020).

73. S. R. Heidemann, R. E. Buxbaum, Tension as a regulator and integrator of axonal growth. Cell Motil. Cytoskeleton 17, 6–10 (1990).

74. J. Nie, L. Guo, K. Li, Y. Wang, G. Chen, L. Li, H. Chen, F. Deng, X. Jiang, T. Zhang, L. Huang, C. Faraco, D. Zhang, C. Guo, P.-T. Yap, X. Hu, G. Li, J. Lv, Y. Yuan, D. Zhu, J. Han, D. Sabatinelli, Q. Zhao, L. S. Miller, B. Xu, P. Shen, S. Platt, D. Shen, X. Hu, T. Liu, Axonal Fiber Terminations Concentrate on Gyri. Cerebral Cortex 22, 2831–2839 (2012).

75. F. Ge, X. Li, M. J. Razavi, H. Chen, T. Zhang, S. Zhang, L. Guo, X. Hu, X. Wang, T. Liu, Denser Growing Fiber Connections Induce 3-hinge Gyral Folding. Cerebral Cortex 28, 1064–1075 (2018).

76. H. Oliveri, K. Franze, A. Goriely, Theory for Durotactic Axon Guidance. Phys. Rev. Lett. 126, 118101 (2021).

77. A. Jakab, E. Schwartz, G. Kasprian, G. M. Gruber, D. Prayer, V. Schapf, G. Langs, Fetal functional imaging portrays heterogeneous development of emerging human brain networks. Front. Hum. Neurosci. 8 (2014).

78. C. Raybaud, T. Ahmad, N. Rastegar, M. Shroff, M. Al Nassar, The premature brain: developmental and lesional anatomy. Neuroradiology 55, 23–40 (2013).

79. L. Vasung, C. Lepage, M. Radoš, M. Pletikos, J. S. Goldman, J. Richiardi, M. Raguž, E. Fischi-Gómez, S. Karama, P. S. Huppi, A. C. Evans, I. Kostovic, Quantitative and Qualitative Analysis of Transient Fetal Compartments during Prenatal Human Brain Development. Front Neuroanat 10, 11 (2016).

80. M. Ouyang, J. Dubois, Q. Yu, P. Mukherjee, H. Huang, Delineation of early brain development from fetuses to infants with diffusion MRI and beyond. NeuroImage 185, 836–850 (2019).

81. L. Vasung, A. Rezayev, H. J. Yun, J. W. Song, A. van der Kouwe, N. Stewart, A. Palani, T. Shiohama, F. Chouinard-Decorte, J. Levman, E. Takahashi, Structural and Diffusion MRI Analyses With Histological Observations in Patients With Lissencephaly. Front. Cell Dev. Biol. 7, 124 (2019).

82. A. Ortug, B. Valli, J. L. Alatorre Warren, T. Shiohama, A. van der Kouwe, E. Takahashi, Brain Pathways in LIS1-Associated Lissencephaly Revealed by Diffusion MRI Tractography. Brain Sci 13, 1655 (2023).

83. C. E. Kelly, D. K. Thompson, C. L. Adamson, G. Ball, T. Dhollander, R. Beare, L. G. Matthews, B. Alexander, J. L. Y. Cheong, L. W. Doyle, P. J. Anderson, T. E. Inder, Cortical growth from infancy to adolescence in preterm and term-born children. Brain 147, 1526–1538 (2024).

84. L. S. Gorham, A. R. Latham, D. Alexopoulos, J. K. Kenley, E. Iannopollo, R. E. Lean, D. Loseille, T. A. Smyser, J. J. Neil, C. E. Rogers, C. D. Smyser, K. Garcia, Children born very preterm experience altered cortical expansion over the first decade of life. Brain Communications 6, fcae318 (2024).

85. Y. H. Jang, H. Kim, J. Y. Lee, J.-H. Ahn, A. W. Chung, H. J. Lee, Altered development of structural MRI connectome hubs at near-term age in very and moderately preterm infants. Cerebral Cortex 33, 5507–5523 (2023).

86. D. J. Ardesch, L. H. Scholtens, S. C. De Lange, L. Roumazeilles, A. A. Khrapitchev, T. M. Preuss, J. K. Rilling, R. B. Mars, M. P. Van Den Heuvel, Scaling Principles of White Matter Connectivity in the Human and Nonhuman Primate Brain. Cerebral Cortex 32, 2831–2842 (2022).

87. M. Ercsey-Ravasz, N. T. Markov, C. Lamy, D. C. Van Essen, K. Knoblauch, Z. Toroczkai, H. Kennedy, A Predictive Network Model of Cerebral Cortical Connectivity Based on a Distance Rule. Neuron 80, 184–197 (2013).

88. C. Papini, L. Palaniyappan, J. Kroll, S. Froudist-Walsh, R. M. Murray, C. Nosarti, Altered Cortical Gyrification in Adults Who Were Born Very Preterm and Its Associations With Cognition and Mental Health. Biological Psychiatry: Cognitive Neuroscience and Neuroimaging 5, 640–650 (2020).

89. K. Im, M. J. Paldino, A. Poduri, O. Sporns, P. E. Grant, Altered white matter connectivity and network organization in polymicrogyria revealed by individual gyral topology-based analysis. NeuroImage 86, 182–193 (2014).

90. D. Batalle, E. J. Hughes, H. Zhang, J.-D. Tournier, N. Tusor, P. Aljabar, L. Wali, D. C. Alexander, J. V. Hajnal, C. Nosarti, A. D. Edwards, S. J. Counsell, Early development of structural networks and the impact of prematurity on brain connectivity. Neuroimage 149, 379–392 (2017).

91. J. Weickenmeier, R. de Rooij, S. Budday, P. Steinmann, T. C. Ovaert, E. Kuhl, Brain stiffness increases with myelin content. Acta Biomaterialia 42, 265–272 (2016).

92. S. Budday, P. Steinmann, E. Kuhl, Physical biology of human brain development. Front. Cell. Neurosci. 9 (2015).

93. S. Budday, R. Nay, R. de Rooij, P. Steinmann, T. Wyrobek, T. C. Ovaert, E. Kuhl, Mechanical properties of gray and white matter brain tissue by indentation. Journal of the Mechanical Behavior of Biomedical Materials 46, 318–330 (2015).

94. K. Miller, Ed., Biomechanics of the Brain (Springer New York, New York, NY, 2011; https://link.springer.com/10.1007/978-1-4419-9997-9)*Biological and Medical Physics, Biomedical Engineering*.

95. K. S. Rockland, J. DeFelipe, Cortical White Matter: Beyond the Pale. Front. Neuroanat. 5 (2012).

96. P. Chavoshnejad, G. K. German, M. J. Razavi, Hyperelastic material properties of axonal fibers in brain white matter. Brain Multiphysics 2, 100035 (2021).

97. H. Hoursan, F. Farahmand, M. T. Ahmadian, A Three-Dimensional Statistical Volume Element for Histology Informed Micromechanical Modeling of Brain White Matter. Ann Biomed Eng 48, 1337–1353 (2020).

98. S. A. Yousefsani, F. Farahmand, A. Shamloo, A three-dimensional micromechanical model of brain white matter with histology-informed probabilistic distribution of axonal fibers. Journal of the Mechanical Behavior of Biomedical Materials 88, 288–295 (2018).

99. K. B. Arbogast, D. F. Meaney, Biomechanical characterization of the constitutive relationship of the brainstem. In: Proceedings of the Society of Automotive Engineers, 153–9 (1995).

100. M. J. Razavi, T. Liu, X. Wang, Mechanism Exploration of 3-Hinge Gyral Formation and Pattern Recognition. Cerebral Cortex Communications 2, tgab044 (2021).

101. M. J. Razavi, T. Zhang, H. Chen, Y. Li, S. Platt, Y. Zhao, L. Guo, X. Hu, X. Wang, T. Liu, Radial Structure Scaffolds Convolution Patterns of Developing Cerebral Cortex. Front. Comput. Neurosci. 11, 76 (2017).

102. K. Schilling, Y. Gao, V. Janve, I. Stepniewska, B. A. Landman, A. W. Anderson, Confirmation of a gyral bias in diffusion MRI fiber tractography. Human Brain Mapping 39, 1449–1466 (2018).

103. M. Cao, Y. He, Z. Dai, X. Liao, T. Jeon, M. Ouyang, L. Chalak, Y. Bi, N. Rollins, Q. Dong, H. Huang, Early Development of Functional Network Segregation Revealed by Connectomic Analysis of the Preterm Human Brain. *Cereb. Cortex*, bhw038 (2016).

104. F. Rheault, P. Poulin, A. Valcourt Caron, E. St-Onge, M. Descoteaux, Common misconceptions, hidden biases and modern challenges of dMRI tractography. J. Neural Eng. 17, 011001 (2020).

105. C. Delettre, A. Messé, L.-A. Dell, O. Foubet, K. Heuer, B. Larrat, S. Meriaux, J.-F. Mangin, I. Reillo, C. De Juan Romero, V. Borrell, R. Toro, C. C. Hilgetag, Comparison between diffusion MRI tractography and histological tract-tracing of cortico-cortical structural connectivity in the ferret brain. Network Neuroscience 3, 1038–1050 (2019).

106. A. Yendiki, M. Aggarwal, M. Axer, A. F. D. Howard, A.-M. V. C. Van Walsum, S. N. Haber, Post mortem mapping of connectional anatomy for the validation of diffusion MRI. NeuroImage 256, 119146 (2022).

107. G. Girard, R. Caminiti, A. Battaglia-Mayer, E. St-Onge, K. S. Ambrosen, S. F. Eskildsen, K. Krug, T. B. Dyrby, M. Descoteaux, J.-P. Thiran, G. M. Innocenti, On the cortical connectivity in the macaque brain: A comparison of diffusion tractography and histological tracing data. NeuroImage 221, 117201 (2020).

108. E. K. Rodriguez, A. Hoger, A. D. McCulloch, Stress-dependent finite growth in soft elastic tissues. Journal of Biomechanics 27, 455–467 (1994).

109. M. Liu, C. Lepage, S. Y. Kim, S. Jeon, S. H. Kim, J. P. Simon, N. Tanaka, S. Yuan, T. Islam, B. Peng, K. Arutyunyan, W. Surento, J. Kim, N. Jahanshad, M. A. Styner, A. W. Toga, A. J. Barkovich, D. Xu, A. C. Evans, H. Kim, Robust Cortical Thickness Morphometry of Neonatal Brain and Systematic Evaluation Using Multi-Site MRI Datasets. Front. Neurosci. 15, 650082 (2021).

110. G. Ball, S. Oldham, V. Kyriakopoulou, L. Z. J. Williams, V. Karolis, A. Price, J. Hutter, M. L. Seal, A. Alexander-Bloch, J. V. Hajnal, A. D. Edwards, E. C. Robinson, J. Seidlitz, Molecular signatures of cortical expansion in the human foetal brain. Nat Commun 15, 9685 (2024).

111. L. Ronan, N. Voets, C. Rua, A. Alexander-Bloch, M. Hough, C. Mackay, T. J. Crow, A. James, J. N. Giedd, P. C. Fletcher, Differential Tangential Expansion as a Mechanism for Cortical Gyrification. Cerebral Cortex 24, 2219–2228 (2014).

112. A. Menzel, Modelling of anisotropic growth in biological tissues. A new approach and computational aspects. Biomech Model Mechanobiol 3, 147–171 (2005).

113. I. Yurchenko, J. M. Vensi Basso, V. S. Syrotenko, C. Staii, Anomalous diffusion for neuronal growth on surfaces with controlled geometries. PLoS ONE 14, e0216181 (2019).

114. S. De Vincentiis, A. Falconieri, M. Mainardi, V. Cappello, V. Scribano, R. Bizzarri, B. Storti, L. Dente, M. Costa, V. Raffa, Extremely Low Forces Induce Extreme Axon Growth. J. Neurosci. 40, 4997–5007 (2020).

115. S. Budday, T. C. Ovaert, G. A. Holzapfel, P. Steinmann, E. Kuhl, Fifty Shades of Brain: A Review on the Mechanical Testing and Modeling of Brain Tissue. Arch Computat Methods Eng, doi: 10.1007/s11831-019-09352-w (2019).

116. M. Cottaar, M. Bastiani, N. Boddu, M. F. Glasser, S. Haber, D. C. Van Essen, S. N. Sotiropoulos, S. Jbabdi, Modelling white matter in gyral blades as a continuous vector field. NeuroImage 227, 117693 (2021).

117. D. C. Van Essen, S. M. Smith, D. M. Barch, T. E. J. Behrens, E. Yacoub, K. Ugurbil, WU-Minn HCP Consortium, The WU-Minn Human Connectome Project: an overview. Neuroimage 80, 62–79 (2013).

118. J.-D. Tournier, R. Smith, D. Raffelt, R. Tabbara, T. Dhollander, M. Pietsch, D. Christiaens, B. Jeurissen, C.-H. Yeh, A. Connelly, MRtrix3: A fast, flexible and open software framework for medical image processing and visualisation. NeuroImage 202, 116137 (2019).

119. B. B. Avants, N. Tustison, G. Song, Advanced normalization tools (ANTS). Insight j 2 2, 1–35 (2009).

120. C. Walter, R. Balouchzadeh, K. E. Garcia, C. D. Kroenke, A. Pathak, P. V. Bayly, Multi-scale measurement of stiffness in the developing ferret brain. Sci Rep 13, 20583 (2023).

121. T. J. Nowakowski, A. Bhaduri, A. A. Pollen, B. Alvarado, M. A. Mostajo-Radji, E. Di Lullo, M. Haeussler, C. Sandoval-Espinosa, S. J. Liu, D. Velmeshev, J. R. Ounadjela, J. Shuga, X. Wang, D. A. Lim, J. A. West, A. A. Leyrat, W. J. Kent, A. R. Kriegstein, Spatiotemporal gene expression trajectories reveal developmental hierarchies of the human cortex. Science 358, 1318–1323 (2017).

122. M. Á. Martínez-Martínez, C. De Juan Romero, V. Fernández, A. Cárdenas, M. Götz, V. Borrell, A restricted period for formation of outer subventricular zone defined by Cdh1 and Trnp1 levels. Nat Commun 7, 11812 (2016).

123. X. Xu, R. Chen, T. Zheng, Z. Zhao, M. Li, D. Wu, Molecular and Cellular Underpinnings of Spatial Heterogeneity in Fetal Cortical Folding. Neuroscience [Preprint] (2026). 10.64898/2026.01.20.700501.

124. T. J. Dennerll, P. Lamoureux, R. E. Buxbaum, S. R. Heidemann, The cytomechanics of axonal elongation and retraction. The Journal of Cell Biology 109, 3073–3083 (1989).

125. D. Bray, Axonal growth in response to experimentally applied mechanical tension. Developmental Biology 102, 379–389 (1984).

126. P. Lamoureux, S. R. Heidemann, N. R. Martzke, K. E. Miller, Growth and elongation within and along the axon. Developmental Neurobiology 70, 135–149 (2010).

127. S. Chada, P. Lamoureux, R. E. Buxbaum, S. R. Heidemann, Cytomechanics of neurite outgrowth from chick brain neurons. Journal of Cell Science 110, 1179–1186 (1997).

128. Y.-J. Chang, C.-J. Tsai, F.-G. Tseng, T.-J. Chen, T.-W. Wang, Micropatterned stretching system for the investigation of mechanical tension on neural stem cells behavior. *Nanomedicine: Nanotechnology*, Biology and Medicine 9, 345–355 (2013).

129. M. J. Razavi, T. Zhang, X. Li, T. Liu, X. Wang, Role of mechanical factors in cortical folding development. *Phys*. Rev. E 92, 032701 (2015).

130. A. Solhtalab, A. H. Foroughi, L. Pierotich, M. J. Razavi, Stress Landscape of Folding Brain Serves as a Map for Axonal Pathfinding. [Preprint] (2024). 10.21203/rs.3.rs-5111417/v1.

131. H. Oliveri, K. Franze, A. Goriely, Theory for Durotactic Axon Guidance. Phys. Rev. Lett. 126, 118101 (2021).

132. J. P. Sunnerberg, M. Descoteaux, D. L. Kaplan, C. Staii, Axonal growth on surfaces with periodic geometrical patterns. PLoS ONE 16, e0257659 (2021).

133. S. Saeidi, M. P. Kainz, M. Dalbosco, M. Terzano, G. A. Holzapfel, Histology-informed multiscale modeling of human brain white matter. Sci Rep 13, 19641 (2023).

134. J. Veraart, D. S. Novikov, D. Christiaens, B. Ades-aron, J. Sijbers, E. Fieremans, Denoising of diffusion MRI using random matrix theory. NeuroImage 142, 394–406 (2016).

135. J. Tournier, F. Calamante, A. Connelly, MRtrix: Diffusion tractography in crossing fiber regions. Int J Imaging Syst Tech 22, 53–66 (2012).

136. R. E. Smith, J.-D. Tournier, F. Calamante, A. Connelly, Anatomically-constrained tractography: Improved diffusion MRI streamlines tractography through effective use of anatomical information. NeuroImage 62, 1924–1938 (2012).

137. B. Jeurissen, J.-D. Tournier, T. Dhollander, A. Connelly, J. Sijbers, Multi-tissue constrained spherical deconvolution for improved analysis of multi-shell diffusion MRI data. NeuroImage 103, 411–426 (2014).

138. J.-D. Tournier, F. Calamante, A. Connelly, Robust determination of the fibre orientation distribution in diffusion MRI: Non-negativity constrained super-resolved spherical deconvolution. NeuroImage 35, 1459–1472 (2007).

139. B. Fischl, FreeSurfer. NeuroImage 62, 774–781 (2012).

