## Supplementary Materials for "A Mechanical Theory for the Formation of Short Association Fibers in the Brain"

**Stabilization and sensitivity analysis of fiber-based connectivity metrics**

To quantitatively assess the proposed stress-dependent reorientation mechanism, we analyzed the temporal evolution of the alignment angle between fiber trajectories and the local MTPS direction (Fig. S1), distinguishing between pre-folding and late-folding initiation regimes based on the bifurcation time $T_\beta$. The results show that alignment evolves in a strongly time-dependent manner and is governed by the timing of fiber growth relative to folding. Fibers initiating prior to bifurcation exhibit low initial misalignment, followed by a pronounced increase near $T_\beta$ due to rapid changes in the stress field, and a subsequent gradual decrease as fibers progressively reorient toward MTPS directions, indicating a delayed but continuous mechanical adaptation. In contrast, fibers initiating after bifurcation display relatively stable alignment with lower variability and no sharp transition, reflecting growth within an already established stress field. Together, these findings demonstrate that fiber reorientation is a gradual process driven by the evolving mechanical environment and critically dependent on the timing of folding.

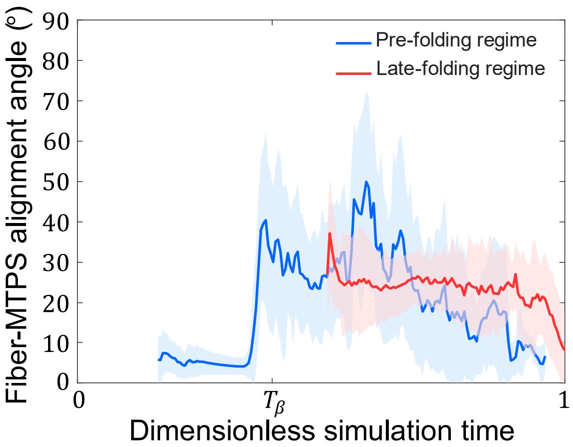


**Fig. S1. Stress-dependent fiber reorientation and frequency of projection-like fibers across different initiation regimes.** The figure shows the temporal evolution of the mean fiber–MTPS alignment angle (solid lines) as a function of pseudo-time $T$, with shaded regions indicating the standard deviation across fibers. Two growth regimes are considered based on the bifurcation time $T_\beta$ (see Fig. 3): pre-folding regime ($T_{init} \leq \frac{2}{3}T_\beta$, blue) and late-folding regime ($T_{init} \geq \frac{4}{3}T_\beta$, red). In the early growth regime, the alignment angle remains low prior to bifurcation, followed by a sharp increase near $T_\beta$, reflecting a transient mismatch between fiber orientation and the rapidly evolving MTPS field. At later times, the angle decreases, indicating gradual reorientation of fibers toward MTPS directions. In contrast, fibers initiating after bifurcation exhibit a more stable alignment behavior with reduced variability, consistent with growth within a relatively stationary stress field. The parameters used in this figure are as follows: $\mu_\mathrm{f}/\mu_\mathrm{s} = 2$, $\mu_\mathrm{c}/\mu_\mathrm{s} = 2$, $a = 0.015\ \mathrm{mm\ Pa^{-1}\ d^{-1}}$, $G^\mathrm{axn} = 0.6\ \mathrm{mm\ d^{-1}}$, where $\mu_\mathrm{f}$, $\mu_\mathrm{s}$, and $\mu_\mathrm{c}$ are the shear moduli of the fiber, subplate, and cortex, respectively, $a$ is the stress-dependent elongation rate, and $G^\mathrm{axn}$ is the axon growth rate.

**Independence of SAFs Characteristics from afferent fiber inclusion**

Our results indicate that the inclusion of afferent fibers does not influence the behavior or developmental characteristics of SAFs (Fig. S2). Therefore, in all models, the growth of afferent fibers was suppressed to isolate the effects of SAFs.

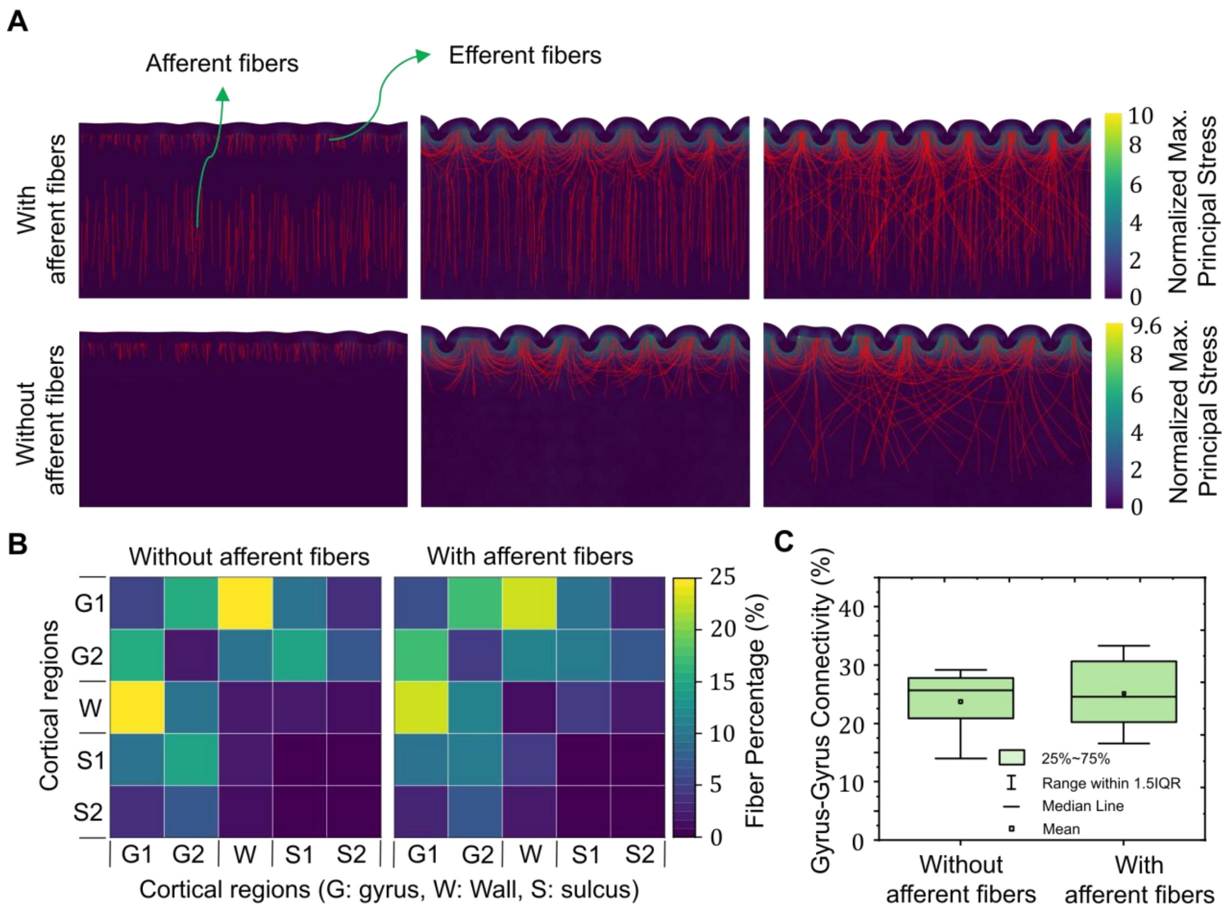


**Fig. S2. Impact of afferent fibers on the behavior and developmental characteristics of short association fibers (SAFs)**. (**A**) Comparison of the dynamic growth and evolution of afferent and efferent fibers during cortical folding for an axon growth rate of $0.8\ \mathrm{mm\ d^{-1}}$. The figure contrasts two scenarios: one with afferent fibers and one without (where fibers grow radially from the core toward the cortex). The parameters used in this analysis are: $\mu_\mathrm{f}/\mu_\mathrm{s} = 2$, $\mu_\mathrm{c}/\mu_\mathrm{s} = 2$, and $a = 0.015\ \mathrm{mm\ Pa^{-1}\ d^{-1}}$, where $\mu_\mathrm{f}$, $\mu_\mathrm{s}$, and $\mu_\mathrm{c}$ are the shear moduli of the fiber, subplate, and cortex, respectively, and $a$ is the stress-dependent elongation rate. (**B**) Connectivity distribution across the gyral span, comparing models with and without afferent fibers. Rows and columns correspond to the same set of regions (G1, G2: gyri; W: wall; S1, S2: sulci), forming a symmetric matrix that represents pairwise connectivity between regions. The regional definitions follow the division of the cortical plate into five equal-area bins from sulci to gyri, as described in Fig. 6. (**C**) Percentage of gyrus-to-gyrus connections for the two models, showing that the inclusion of afferent fibers does not affect SAFs or their connectivity patterns. The box plots illustrate the interquartile range (IQR), extending from the 25th to the 75th percentile. The mean is indicated by a white dot, the median by a black line within the box, and whiskers represent data within 1.5 times the IQR.

### Computational timing regimes and qualitative biological context for SAF development

Table S1 presents two complementary but independently defined descriptions of SAF organization. Panel A summarizes the computational timing regimes, which are defined relative to the mechanically determined bifurcation time, $T_\beta$. Panel B summarizes the progression of SAF morphology reported in fetal imaging studies between approximately 22 and 38 gestational weeks (GW) (*52*). The reported biological progression motivated the qualitative structure and ordering of the computational regimes; however, the numerical regime boundaries were defined relative to $T_\beta$ and were not obtained by converting model time into gestational age.

Table S1. Computational timing regimes and independently reported fetal SAF observations

| **Panel A. Computational timing regimes** | | | |
|---|---|---|---|
| **Computational timing regime** | **Definition relative to $T_\beta$** | **Simulated fiber organization** | **Folding-related mechanical environment** |
| Pre-folding regime | $T_{init} \leq \frac{2}{3}T_\beta$ | Predominantly long, projection-like trajectories; only a small subset of fibers forms U-shaped connections | Folding-induced stress field is not yet established; fiber trajectories are primarily influenced by initial orientation and stochastic tip dynamics |
| Peri-folding onset regime | $\frac{2}{3}T_\beta \leq T_{init} \leq T_\beta$ | Mixed organization consisting of projection-like fibers and early-stage U-fibers; initial directional bias begins to emerge | Evolving folding-induced stress gradients begin to influence fiber trajectories |
| Active-folding regime | $T_\beta \leq T_{init} \leq \frac{4}{3}T_\beta$ | Substantial fiber reorientation and increased formation of localized U-fibers connecting adjacent gyri | Strong coupling between the developing folding-induced stress field and fiber growth; stress-dependent reorientation becomes dominant |
| Late-folding / post-folding regime | $\frac{4}{3}T_\beta \leq T_{init}$ | Predominantly localized, well-defined U-shaped trajectories connecting neighboring gyri | Folding geometry and the associated stress landscape are substantially established and strongly constrain fiber trajectories |
| **Panel B. Independently reported fetal observations** | | | |
| **Gestational interval** | **Developmental context** | **SAF morphology** | **Folding context** |
| Approximately 22–26 GW | Early development with limited cortical folding | Relatively flat, loosely organized, and multidirectional superficial pathways without clearly defined U-shaped organization | Limited influence of cortical folding on fiber organization |
| Approximately 26–30 GW | Folding initiation and early developmental transition | SAFs remain relatively dispersed but begin to exhibit partial organization and directional bias | Emerging cortical folds increasingly influence the organization of superficial pathways |
| Approximately 30–34 GW | Rapid folding and structural reorganization | SAFs undergo substantial reorientation and increasingly develop curved trajectories associated with the folding geometry | Increasing folding complexity is accompanied by pronounced reorganization of SAF trajectories |
| Approximately 34–38 GW and later | Late gestational organization and refinement | Increasingly organized U-shaped association bundles are observed between neighboring gyri | Cortical folding geometry is substantially established |

* The gestational intervals in Panel B indicate developmental stages at which the respective SAF morphologies have been reported in fetal observations (*52*). Panels A and B are independently defined: no value of $t$, $T$, $T_{init}$, or $T_\beta$ is assigned to a specific gestational week. The 0–50-day computational interval and its normalized representation are not equivalent to the 22–38 GW observational interval, and the gestational ranges were not used to determine the parameters in Table 1. The biological observations informed the qualitative ordering of the computational regimes, whereas the fractional cutoffs remain operational modeling thresholds. Accordingly, the comparison is limited to shared morphological features and developmental ordering and does not constitute a one-to-one temporal mapping or a prediction of exact gestational transitions.

**Effect of biomechanical parameters on SAF patterning**

*Effect of relative growth rate of cortex and axons.* We investigated the influence of the relative growth rates between the cortical plate and fibers on the development, patterning, and connectivity of SAFs in the folding brain and developing white matter. To modulate this relative growth, we varied the fiber growth rate while keeping cortical expansion constant. An increase in fiber growth rate can also be interpreted as a relative reduction in cortical plate growth, thereby altering the timing and spatial interplay between folding and fiber development.

Figure S3A illustrates the growth and development of SAFs with different growth rates in a system undergoing cortical folding. Simulations were performed using four distinct axonal growth rates $G^{\mathrm{axn}} = 0.6$, $0.8$, $1.0$, and $1.2\ \mathrm{mm\,d^{-1}}$, as summarized in Table 1. Across all scenarios, the stress-dependent elongation rate was held constant at $a = 0.015\ \mathrm{mm\,Pa^{-1}\,d^{-1}}$, along with all other simulation parameters. The simulations reveal that increasing the axonal growth rate from $0.6$ to $1.2\ \mathrm{mm\,d^{-1}}$ reduces the number of SAFs while increases the number of long projection fibers. This behavior arises because, at higher fiber growth rates or lower cortical plate growth rates, fibers originating near the cortical boundary quickly traverse the high-stress regions adjacent to the cortex and extend into the lower-stress regions of the deep subplate without experiencing significant stress-dependent reorientation. In the lower-stress regions, tensile stress exerts a minimal influence on fiber reorientation. As a result, most fibers grow in a nearly straight trajectory after leaving the high-stress regions, forming long projection fibers. It is interesting to observe that, similar to the brain, in addition to the predominantly U-shaped fibers that connect adjacent gyri, longer SAFs also form connections with more distant gyri, extending to the second or third gyrus beyond the immediately adjacent one. It is shown that a higher axonal growth rate alters the association time between folding and connectivity development, shifting it toward the pre-folding regime, as discussed in Fig. 3 of the main text. Under these conditions, most fibers develop into long projection fibers, while a smaller subset forms U-fibers.

Figure S3B presents the distribution of SAFs connections along the gyral span for the different growth rates. The connectivity matrices indicate that for $G^{\mathrm{axn}} = 0.6\ \mathrm{mm\,d^{-1}}$, streamlines connecting gyri to other gyri (either the same or different) are the most prevalent. These are followed by streamlines connecting gyri to walls and gyri to sulci, while sulcus-to-sulcus connections remain the least common. These observations are consistent with imaging data that highlight the prevalence of gyrus-to-gyrus connectivity within SAFs(*1*). As the growth rate increases, the prevalence of gyrus-to-gyrus connections decreases, while the proportion of streamlines connecting gyri to walls increases, with sulcus-to-sulcus connections remaining consistently rare. Fig. S3C highlights the effect of axonal growth rate on gyrus-to-gyrus connectivity. It shows that as the growth rate increases, the percentage of streamlines forming gyrus-to-gyrus connections declines. A significant correlation is observed between the connectivity patterns at $G^{\mathrm{axn}} = 0.6\ \mathrm{mm\,d^{-1}}$ and other growth rates.

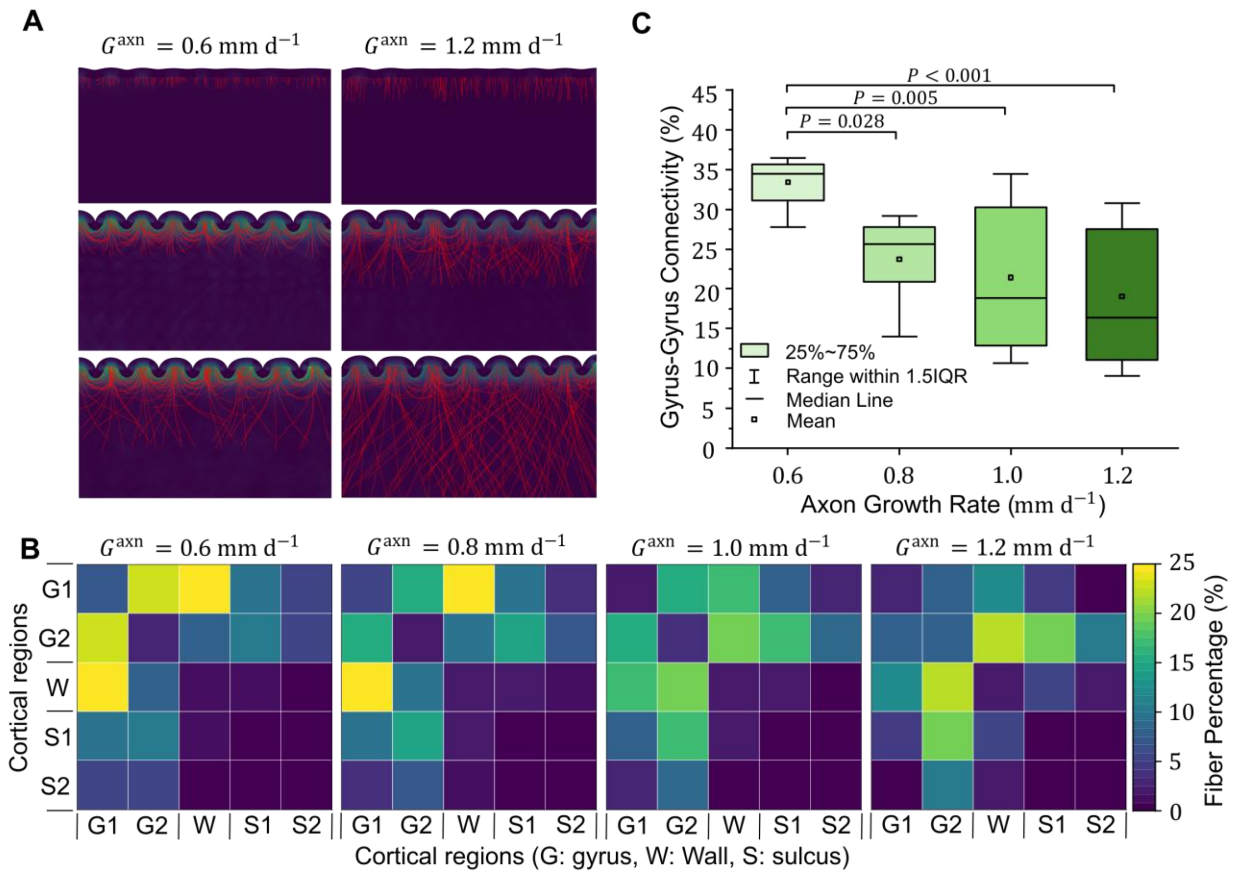


**Fig. S3. Axonal growth rate influences the development and organization of short association fibers (SAFs).** (**A**) dynamic growth and evolution of SAFs and long projection fibers for axonal growth rates $0.6\ \mathrm{mm\ d^{-1}}$ and $1.2\ \mathrm{mm\ d^{-1}}$ during cortical folding. The used parameters in this figure are: $\mu_{\mathrm{f}}/\mu_{\mathrm{s}} = 2$, $\mu_{\mathrm{c}}/\mu_{\mathrm{s}} = 2$, and $a = 0.015\ \mathrm{mm\ Pa^{-1}\ d^{-1}}$, where $\mu_{\mathrm{f}}$, $\mu_{\mathrm{s}}$, and $\mu_{\mathrm{c}}$ are the shear moduli of the fiber, subplate, and cortex, respectively, and $a$ is the stress-dependent elongation rate. (**B**) connectivity distribution across the gyral span for different axonal growth rates. Rows and columns correspond to the same set of regions (G1, G2: gyri; W: cortical wall; S1, S2: sulci), forming a symmetric matrix that represents pairwise connectivity between regions. The regional definitions follow the division of the cortical plate into five equal-area bins from sulci to gyri, as described in Fig. 6. (**C**) dependency of the percentage of gyrus-to-gyrus connections to the growth rate of fibers, illustrating a significant reduction with higher growth rates. The box plots represent the interquartile range (IQR), extending from the 25th to the 75th percentile. The mean is depicted as a white dot, while the median is shown as a black line within the box. The whiskers indicate the data range within 1.5 times the IQR.

*Effect of cortex stiffness on SAFs development and patterning.* We also investigated how the stiffness ratio between the cortex and the subplate ($\mu_c/\mu_s$) influences the navigation of efferent fibers, the formation of SAFs within the subplate, and their connectivity. Four distinct stiffness ratios (1, 2, 3, and 4) were examined, keeping all other parameters constant across the models. This range was chosen for the parametric study because the stiffness ratio between the cortex and subplate in a developing brain can vary significantly due to dynamic changes in brain tissue properties during cortical folding(*120*). However, experimental evidence from mature brains suggests that cortical stiffness is typically 1 to 2 times that of the white matter(*59*, *93*, *120*).

Figure S4A-D presents the simultaneous cortical folding and SAFs development observed across models with four different stiffness ratios. As the stiffness ratio increases, the bifurcation time ($T_\beta$), the onset of distinct morphological changes leading to fold formation, occurs earlier(*30*). Bifurcation marks the point at which the initially smooth cortical surface begins to develop gyri and sulci, coinciding with instability and a rapid increase in the system's dynamic energy. An earlier bifurcation shifts the association timing between folding and connectivity development toward the pre-folding regime, as discussed in Fig. 3. Therefore, the models predict that higher stiffness ratios accelerate the folding process, prompting an earlier onset of stress-dependent fiber reorientation. At higher cortical stiffness ratios (e.g., 4), fibers are more likely to navigate within the elevated stress region near the cortex, reorienting toward the direction of the MTPS (Fig. S4D). This results in the formation of a greater number of SAFs. Conversely, at lower stiffness ratios (e.g., 1), bifurcation occurs later, allowing most fibers to traverse through the high-stress regions before folding begins (Fig. S4A). This temporal difference in folding profoundly influences the final organization of SAFs in the subplate. For a stiffness ratio of 1, the majority of fibers follow long trajectories, connecting distant regions. In contrast, at higher stiffness ratios, a greater proportion of fibers take shorter paths, resulting in an increased number of SAFs and a corresponding decrease in projection fibers. Moreover, when bifurcation is delayed, as seen with a stiffness ratio of 1, most fibers traverse high-stress regions where the MTPS directions form characteristic U-shaped patterns, as depicted in Fig. 2B. Subsequently, these fibers navigate through low-stress regions where the MTPS directions are predominantly vertical. On the other hand, when bifurcation occurs earlier (as with higher stiffness ratios), most fibers undergo reorientation in the U-shaped MTPS regions, resulting in the formation of SAFs.

The quantitative analysis of fiber connectivity, as shown in Fig. S4E-F, underscores the visual observations. Fig. S4E presents the distribution of fiber connections along the gyral span for stiffness ratios of 1 (top) and 4 (bottom). The connectivity matrices indicate that, at a stiffness ratio of 4, streamlines connecting gyri to other gyri (either the same or different) are predominant. Figure S4F shows the percentage of fibers that connect gyrus to gyrus for different stiffness ratios. Statistical analysis reveals a significant correlation between a stiffness ratio of 1 and higher stiffness ratios, highlighting that increasing the stiffness ratio leads to a greater proportion of SAFs connecting adjacent gyri. Figure S4G further illustrates this trend by quantifying the total number of SAFs formed at the final stage of cortical folding ($T = 1$) across varying stiffness ratios. As the stiffness ratio increases, a notable rise in the number of SAFs is observed. This suggests that a stiffer cortex provides a more favorable mechanical environment for the emergence and stabilization of SAFs, particularly during the later stages of cortical development. This observation can be explained by the fact that increased cortical stiffness leads to an earlier onset of folding, allowing more time for the stress field induced by cortical folding to reorient fibers toward MTPS directions. As folding progresses, these stable mechanical cues further guide fiber trajectories, promoting the formation of SAFs that connect adjacent gyri. Overall, these findings underscore the importance of tissue mechanics in guiding the structural organization of white matter connectivity, specifically the development of localized cortico-cortical pathways such as SAFs.

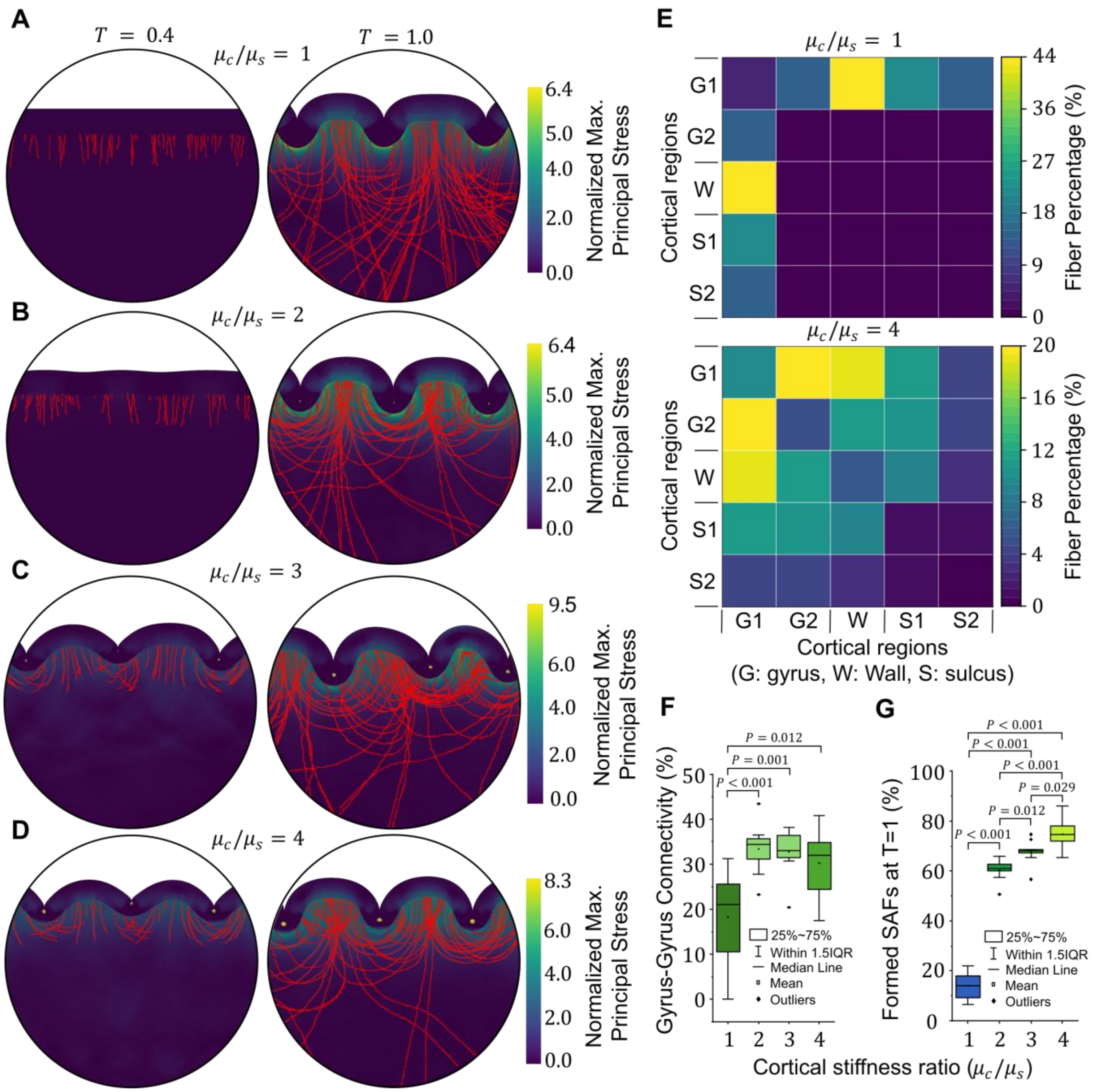


**Fig. S4. Increasing cortical stiffness facilitates the formation and establishment of SAFs.** (**A-D**) Connectivity development and patterning for models with varying stiffness ratio of the cortex to the subplate: (A) $\mu_c/\mu_s = 1$, (B) $\mu_c/\mu_s = 2$, (C) $\mu_c/\mu_s = 3$ ,and (D) $\mu_c/\mu_s = 4$. As the stiffness ratio increases, bifurcation time ($T_\beta$) occurs earlier, resulting in faster cortical folding and more rapid initiation of stress-dependent fiber reorientation. At higher stiffness ratios (e.g., $4$), the cortical surface undergoes folding earlier, leading to fibers being reoriented toward the direction of maximum tensile principal stress (MTPS) in the process. In contrast, at lower stiffness ratios (e.g., $1$), bifurcation happens later, allowing fibers to pass through high-stress regions before reorientation begins. The parameters used in this figure are as follows: $\mu_f/\mu_s = 2$, $a = 0.015\ \mathrm{mm\ Pa^{-1}\ d^{-1}}$, $G^{\mathrm{axn}} = 0.6\ \mathrm{mm\ d^{-1}}$, where $\mu_f$ and $\mu_s$ are the shear moduli of the fiber and subplate, respectively, $a$ is the stress-dependent elongation rate, and $G^{\mathrm{axn}}$ is the axonal growth rate. (**E**) Distribution of fiber connections along the gyral span for stiffness ratios of $1$ and $4$. The values are the average of $10$ individual cases. Rows and columns correspond to the same set of regions (G1, G2: gyri; W: wall; S1, S2: sulci), forming a symmetric matrix that represents pairwise connectivity between regions. The regional definitions follow the division of the cortical plate into five equal-area bins from sulci to gyri, as described in

Fig. 6. (**F**) Percentage of fibers connecting gyrus to gyrus for different stiffness ratios, demonstrating a significant correlation between stiffness ratio of 1 and higher ratios. (**G**) Total number of SAFs formed at the final stage of cortical folding ($T = 1$) for different stiffness ratios of cortex to subplate. The results show a positive relationship between cortex-to- subplate stiffness ratio and SAF formation, with higher stiffness ratios leading to a greater number of SAFs. Statistical analyses were two-sided and employed Tukey's post hoc method for multiple comparisons. The box plots represent the interquartile range (IQR), extending from the 25th to the 75th percentile. The mean is depicted as a white dot, while the median is shown as a black line within the box. The whiskers indicate the data range within 1.5 times the IQR.

*Effect of fiber stiffness on fold placement.* Here, the influence of fiber stiffness on cortical folding patterns, with a particular focus on how this stiffness affects the formation of sulci and gyri, is discussed. To investigate this, we examined four fiber-to- subplate stiffness ratios, ranging from 1 to 4, with fiber growth initiated during the peri-folding onset regime (Fig. S5). We intentionally concentrated 30% of the fibers in a specific region near the cortical surface, targeting an area where sulci typically form, for a stiffness ratio of $\mu_f/\mu_s = 1$, which represents homogeneous white matter (subplate+axon) (Fig. S5A). The remaining 70% of the fibers were randomly distributed along the cortex-subplate interface. The 30% concentrated and 70% distributed seeding pattern was introduced as a simplified heterogeneous initiation scenario to examine whether localized enrichment of fiber emergence can bias fold placement. This assumption is qualitatively supported by developmental studies showing that prospective gyral regions exhibit nonuniform progenitor-related programs and increased subventricular zone (SVZ) or outer subventricular zone (oSVZ) activity before or during fold formation(*121–123*). To further evaluate the robustness of this assumption, we conducted a sensitivity analysis of the seeding ratio. The results indicate that the observed behavior is not dependent on the specific concentration ratio; rather, the presence of localized fiber enrichment is sufficient to drive fold localization, consistently leading to the formation of a gyrus at the site of enrichment.

In Fig. S5A, when the stiffness of the fibers and the subplate is the same, the fibers do not contribute to fold placement, and the white matter behaves as a homogeneous substrate for cortical folding, resulting in uniform folds with symmetrical patterns. In this scenario, the locations of gyri and sulci are not dictated by the fibers or their concentrated distribution; rather, their positioning is governed by the geometric and mechanical properties of the cortex and the underlying homogeneous white matter. Nevertheless, the stress field generated by cortical folding still influences the navigation of fibers. Interestingly, we observed that when the fibers are stiffer than the subplate (e.g., $\mu_f/\mu_s \geq 2$), the region with a high concentration of fibers consistently develops into a gyrus following cortical folding. This observation held true for all models with stiffer fibers than the subplate (Fig. S5B-D), regardless of the association time between cortical folding and connectivity development. This finding may explain why fiber denisty in gyri is higher than sulci which will be discussed further later. Regarding the effect of the fiber stiffness ratio on connectivity patterning, visual inspection (not quantified) of the models reveals that increasing fiber stiffness results in the formation of deeper U-shaped fibers. This can be attributed to the fact that stiffer fibers are more resistant to reorientation by the stress field of the subplate compared to softer fibers, allowing them to penetrate further into the subplate before changing their trajectory. The effect of fiber stiffness on fold placement in the complete timeline of the association between cortical folding and connectivity development is presented in Fig. S6.

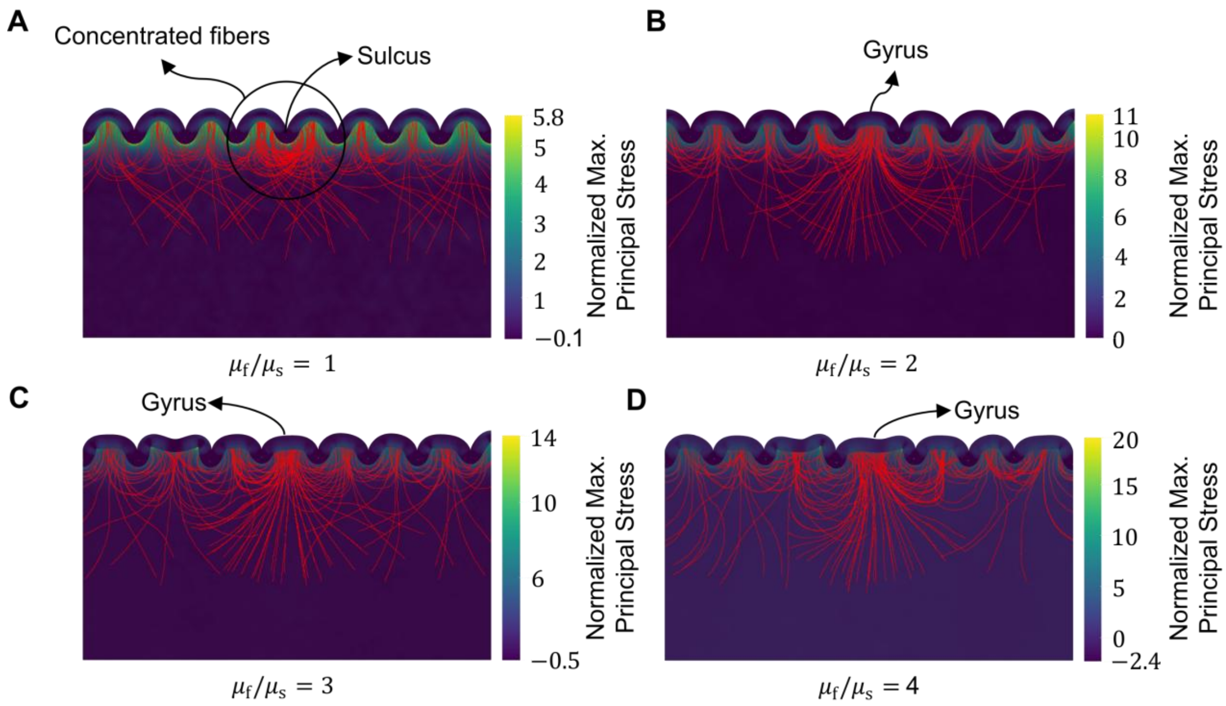


**Fig. S5. Fibers stiffer than the subplate can contribute to the placement of gyri and sulci.** The figure illustrates cortical folding outcomes for four fiber-to-subplate stiffness ratios: (**A**) $\mu_f/\mu_s = 1$, (**B**) $\mu_f/\mu_s = 2$, (**C**) $\mu_f/\mu_s = 3$, and (**D**) $\mu_f/\mu_s = 4$. The results highlight the effect of fiber concentration near the cortex. In the stiffness ratio of $1$, the concentrated region corresponds to sulci, whereas at higher stiffness ratios ($\geq 2$), the same region transitions into a gyrus. The findings demonstrate the influence of mechanical stiffness and fiber distribution on cortical folding dynamics, emphasizing the role of stiffer fibers in promoting gyri formation in high-density regions. The figure is based on the model corresponding to the peri-folding onset regime of fiber growth initiation. The constant parameters used in this figure are: $\mu_c/\mu_s = 2$, $a = 0.015\ \mathrm{mm\ Pa^{-1}\ d^{-1}}$, $G^{\mathrm{axn}} = 0.6\ \mathrm{mm\ d^{-1}}$, where $\mu_s$ and $\mu_c$ are the shear moduli of the subplate and cortex, respectively, $a$ is the stress-dependent elongation rate, and $G^{\mathrm{axn}}$ is the axonal growth rate.

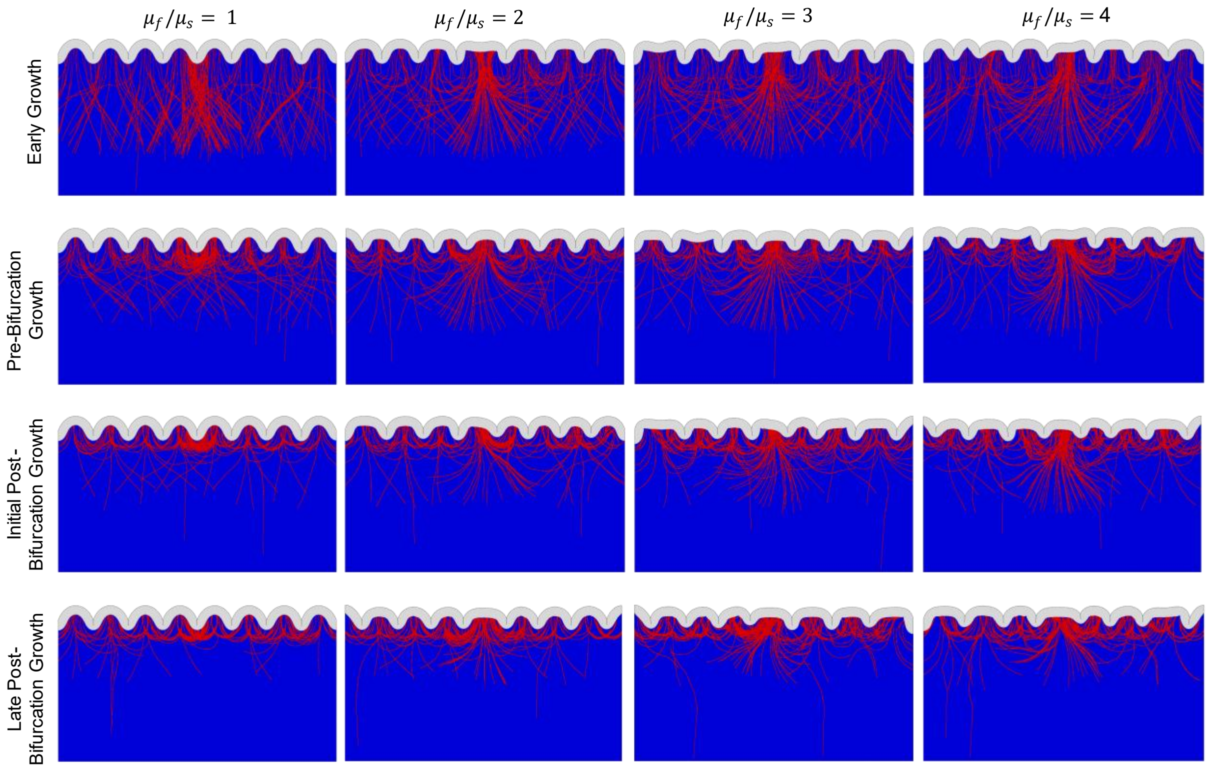


**Fig. S6. Impact of fiber stiffness ratio on cortical folding patterns during the different folding-connctivity association time.** The figure illustrates cortical folding outcomes for four stiffness ratios ($1-4$), highlighting the effect of fiber concentration near the cortex. In the stiffness ratio of 1, the concentrated region corresponds to sulci, whereas at higher stiffness ratios ($\geq 2$), the same region transitions into a gyrus. The findings demonstrate the influence of mechanical stiffness and fiber distribution on cortical folding dynamics, emphasizing the role of stiffer fibers in promoting gyri formation in high-density regions. This trend is reaapiting for all the folding-connctivity association times. The constant parameters used in this figure are: $\mu_c/\mu_s = 2$, $a = 0.015\ \mathrm{mm\ Pa^{-1}\ d^{-1}}$, $G^{\mathrm{axn}} = 0.6\ \mathrm{mm\ d^{-1}}$, where $\mu_s$ and $\mu_c$ are the shear moduli of the subplate and cortex, respectively, $a$ is the stress-dependent elongation rate, and $G^{\mathrm{axn}}$ is the axonal growth rate.

### Methods

**Constitutive framework.** The details of the implemented constitutive framework are presented and discussed herein.

<u>*Discrete model of stress-dependent axon reorientation.*</u> To capture the evolution of fiber directions, we introduce a discrete model to solve Eq. (10) in the main text. We consider a finite time interval $[t_k,\ t_{k+1}]$ which time increment is defined as $\Delta t = t_{k+1} - t_k$. We assume that the preferred direction's vector is known at time $t_k$. First of all, we apply a Gaussian noise to the preferred direction to incorporate stochastic growth dynamics:

$$\mathbf{n}_k^{\mathrm{A}} = \cos(\theta + \epsilon)\,\mathbf{e}_x + \sin(\theta + \epsilon)\,\mathbf{e}_y \tag{S1}$$

where θ is the angle between $\mathbf{n}_k^{\mathrm{A}}$ and the x-axis, $\epsilon$ represents the Gaussian noise, and $\mathbf{e}_x$ and $\mathbf{e}_y$ are the unit vectors in the $x$- and $y$-axes, respectively. Within each time step, the reorientation vector is calculated, and we update $\mathbf{n}_k^{\mathrm{A}}$ iteratively using an exponential mapping scheme:

$$\mathbf{n}_{k+1}^{\mathrm{A}} = \exp(-\Delta t \boldsymbol{\varepsilon} \cdot \boldsymbol{\omega}_k) \cdot \mathbf{n}_k^{\mathrm{A}} = \boldsymbol{R}_k \cdot \mathbf{n}_k^{\mathrm{A}} \tag{S2}$$

where $\mathbf{n}_k^{\mathrm{A}}$ and $\mathbf{n}_{k+1}^{\mathrm{A}}$ are the preferred directions at $t_k$ and $t_{k+1}$, respectively, and $\boldsymbol{\varepsilon}$ is the third-order permutation symbol. The proper orthogonal tensor $\boldsymbol{R}_k$ is defined by the following closed-form expression

$$\boldsymbol{R}_k = \cos(\Delta t \omega_k)\, \mathbf{I} - \sin(\Delta t \omega_k)\, \boldsymbol{\varepsilon} \cdot \mathbf{n}_k^{\omega} + [1 - \cos(\Delta t \omega_k)] \mathbf{n}_k^{\omega} \otimes \mathbf{n}_k^{\omega} \tag{S3}$$

After determining $\mathbf{n}_{k+1}^{\mathrm{A}}$, the rotation vector $\boldsymbol{\omega}_k$ can be updated by approximating $\mathbf{n}_k^{\mathrm{A}} \approx \mathbf{n}_{k+1}^{\mathrm{A}}$ and using Eq. (9) in the main text. As the preferred direction reorients toward the direction of maximum tensile principal stress, the fiber begins to grow along the newly aligned direction, as illustrated in Fig. 6B. This formulation provides a nuanced simulation of fiber growth and alignment in response to mechanical stress.

*Tip and stress-induced axon growth.* Our model incorporates two distinct mechanisms of axon elongation: tip-driven growth and stress-induced stretching. Tip-driven growth or intrinsic growth occurs at the axon's growth cone, where forces generated at the tip propel forward movement. This growth type depends on the availability of biochemical signals and nutrients, allowing elongation to proceed independently of external mechanical forces if these resources are present(*113*). In contrast, stress-induced growth, also known as stretch-dependent or towed growth, arises from mechanical stretching along the axon's shaft due to external forces or deformations in the ECM(*39*). Experimental studies indicate that when axons are subjected to persistent external forces, they undergo gradual length adjustments to maintain a preferred tension level(*124–126*). Notably, axons exposed to minimal tension may even exhibit accelerated growth rates, highlighting the adaptability of axonal growth mechanisms in varying mechanical environments(*114*, *127*). It has been experimentally demonstrated that neurites exhibit intrinsic growth in the absence of external forces, while their growth rate increases under tensile loading and their orientation adjusts toward the direction of the applied force(*128*).

In the model, we incorporate both a baseline (fixed) growth rate to mimic the intrinsic growth and a stress-dependent elongation rate. This combined approach allows us to simulate fiber growth in a specific direction, where each time increment reflects the influence of applied tension alongside an intrinsic growth rate, denoted as $G^{\mathrm{axn}}$. Consequently, the total growth rate of the fiber integrates both the stress-induced component and a fixed growth rate providing a comprehensive view of axonal elongation under varied mechanical influences, as formulated in Eq. (11) in the main text. The parameter $a > 0$ defines the sensitivity of axonal elongation to mechanical stress, effectively controlling the rate of stress-dependent growth. In our simulations, we set $\sigma_0$ to zero, meaning that any non-zero stress promotes elongation. This approach aligns with experimental observations indicating that embryonic neurons, such as those in the forebrain, respond to minimal tension with significant growth, reflecting a low threshold for tension-induced axonal elongation(*114*). The growth rate derived from this equation is used to compute the change in fibers length, $\Delta L$, over each time step $\Delta t$ using the Euler method: $\Delta L = \frac{\partial G}{\partial t} \cdot \Delta t$. The new coordinates of the fiber tip $(x_{k+1}, y_{k+1})$ are determined based on its current position $(x_k, y_k)$ and the polar coordinates $\Delta L$ and $\theta$, where $\theta$ is the angle between the preferred direction $\boldsymbol{n}^A$ and the horizontal axis:

$$\begin{aligned} x_{k+1} &= x_k + \Delta L \cdot \cos(\theta) \\ y_{k+1} &= y_k + \Delta L \cdot \sin(\theta) \end{aligned} \tag{S4}$$

This model provides a dynamic framework for simulating axonal growth, capturing the interplay between mechanical stress and directional preferences. The incorporation of the Euler method allows for the numerical exploration of fiber elongation over discrete time intervals, offering insights into the spatiotemporal aspects of neural development. A summary of the entire algorithm for the reorientation of fiber is presented in Table S1.

| **Table S1**. Algorithm update scheme for the reorientation of fiber (axon bundle) |
|---|
| **1. Initialization:**<br>• Set the initial direction vector as $\mathbf{n}^{\mathrm{A}} = \mathbf{n}_k^{\mathrm{A}} = \left[n_{k1}^{\mathrm{A}},\ n_{k2}^{\mathrm{A}},\ n_{k3}^{\mathrm{A}}\right]^t$.<br>• Compute the stress tensor as $\mathbf{T} = J^{\mathrm{e}^{-1}}\mathbf{F}^{\mathrm{e}}.\frac{\partial W}{\partial \mathbf{F}^{\mathrm{eT}}}$ |
| **2. Compute principal stresses and directions:**<br>• Decompose the stress tensor into principal component: $\mathbf{T} = J^{\mathrm{e}^{-1}}\mathbf{F}^{\mathrm{e}}.\frac{\partial W}{\partial \mathbf{F}^{\mathrm{eT}}} = \sum_{i=1}^{3} \sigma_i \mathbf{n}_i \otimes \mathbf{n}_i$, where $\sigma_1 \geq \sigma_2 \geq \sigma_3$<br>• Set the maximum stress direction $\mathbf{n}_{\max}^T = \mathbf{n}_1^T$. |
| **3. Evaluate conditions for stress and direction:**<br>• If $\sigma_1 < 0$ or $\sigma_1 = \sigma_2$ then<br>$\dot{\mathbf{n}}^{\mathrm{A}} = 0$<br>Else if $\mathbf{n}^{\mathrm{A}} \cdot \mathbf{n}_1^T < 0$ then<br>$\mathbf{n}_{\max}^T \leftarrow -\mathbf{n}_1^T$<br>Else<br>$\mathbf{n}_{\max}^T$ remains as $\mathbf{n}_1^T$<br>End if |
| **4. Caculate rotation velocity:**<br>• Compute the magnitude of the rotation velocity as $\omega = \frac{\pi}{2t^*} \lVert \mathbf{n}^{\mathrm{A}} \times \mathbf{n}_{\max}^T \rVert$<br>• Determine the rotation axis (unit vector) as $\mathbf{n}^{\omega} = \frac{\mathbf{n}^{\mathrm{A}} \times \mathbf{n}_{\max}^T}{\lVert \mathbf{n}^{\mathrm{A}} \times \mathbf{n}_{\max}^T \rVert}$ |
| **5. Update the preferred direction vector:**<br>• Update using the following expression:<br>$\mathbf{n}_{k+1}^{\mathrm{A}} = \cos(\Delta t\omega)\, \mathbf{n}^{\mathrm{A}} + \sin(\Delta t\omega)\, \mathbf{n}^{\omega} \times \mathbf{n}^{\mathrm{A}} + [1 - \cos(\Delta t\omega)][\mathbf{n}^{\omega} \cdot \mathbf{n}^{\mathrm{A}}]\mathbf{n}^{\omega}$<br>• Set the new direction vector as $\mathbf{n}_k^{\mathrm{A}} = \mathbf{n}_{k+1}^{\mathrm{A}}$. |

### Finite element model

In the finite element (FE) model, shown in Fig. 6C, we simulate the developing brain as a simplified bilayer structure, consisting of a tangentially expanding cortical layer (top) and a subcortical layer (bottom) consisting of subplate and growing fibers. The constitutive models described earlier were implemented in the Abaqus/Explicit FE software via a user-defined material subroutine (VUMAT). Cortical thickness in mature brains varies by region, typically between $2\ \mathrm{mm}$ and $4\ \mathrm{mm}$, and in this model, we initialized the cortex thickness, $t_c$, at $1.5\ \mathrm{mm}$ and the subplate layer thickness, $h$, at $30\ \mathrm{mm}$. The model width was set to $w = 60\ \mathrm{mm}$ to accommodate multiple folds within the computational domain while minimizing boundary effects. Across simulations with varying parameters (Table 1), the domain produced approximately 7–9 folds, corresponding to a fold wavelength of about ~5–10 mm(*129*) (measured crest-to-crest). This range is consistent with reported intersulcal distances in the developing human cortex and aligns with prior mechanical models of cortical folding, where the characteristic wavelength is governed primarily by cortical thickness and stiffness contrast. Both layers were meshed using 4-node plane strain quadrilateral elements (CPE4R) with a mesh size of $0.1\ \mathrm{mm}$, chosen based on mesh convergence tests. To drive cortical folding, the cortex was set to grow tangentially along the $x$ −axis (as shown in Fig. 6A) at a rate of $G^{\mathrm{c}} = 0.02\ \mathrm{d}^{-1}$ with a growth ratio of $\frac{G^{\mathrm{c}}}{G^{\mathrm{s}}} = 6$ relative to the sublayer(*130*). This differential growth between the cortex and subcortex serves as the primary driver for cortical expansion and gyrification. Symmetric boundary conditions were applied along the $x$ −axis on the left and right sides, while the base of the model was constrained in the $y$ direction.

Each element within the subcortical layer represents either a subplate component or a fiber, each with distinct mechanical properties (Fig. 6A). As shown in Fig. 6C, we model fiber growth and dynamic directional pathfinding by considering two types of fibers: (a) afferent fibers, which originate deep within the subplate and extend upward toward the cortex, forming long-range connections between cortical and subcortical regions, and (b) efferent fibers that emerge near the cortex-subplate interface and grow toward the deep subplate. This framework captures the formation and development of both descending and ascending fibers.

Individual fibers were initialized at random position, either near the cortical layer or deep within the subplate, within an $x$ −range of $0 - 60\ \mathrm{mm}$. The initial growth angles were randomly assigned: $\theta = \frac{\pi}{2} \pm 10\%$ for fibers originating deep within the subplate and $\theta = \frac{3\pi}{2} \pm 10\%$ for those originating near the cortex. The preferred growth direction, $\boldsymbol{n}^A = \cos(\theta + \epsilon)\mathbf{e}_x + \sin(\theta + \epsilon)\,\mathbf{e}_y$, where $\mathbf{e}_x$ and $\mathbf{e}_y$ are unit vectors along the $x -$ and $y$ −axes, respectively, was stored locally at each integration point as an internal variable. The stochastic term $\epsilon$ represents variability in axonal growth direction due to non-mechanical factors such as biochemical cues. It is implemented as additive angular noise sampled from a Gaussian distribution with zero mean and a standard deviation of 0.025 rad(*131*). This introduces small directional perturbations while preserving the dominant influence of stress-dependent growth. During each time step $\Delta t$, the preferred direction $\boldsymbol{n}^A$ was reoriented along the direction of MTPS, following Eq. (S2), allowing the fiber to growth in the updated direction. The fiber's tip position was recalculated at each iteration using Eq. (S4), and the subplate element containing the updated tip position was reclassified as a fiber element. This iterative process continued until the final time step, with each element classified as either a fiber or subplate element, as illustrated in Fig. 6A. The relaxation parameter $t^*$ was selected based on a sensitivity analysis (Fig. S7). We explored a range of values and constrained its selection by requiring smooth and biologically plausible fiber trajectories consistent with gradual axonal reorientation observed experimentally. Small $t^*$ values lead to overly rapid and non-physical reorientation, while large $t^*$ values reduce the ability of fibers to align with the evolving stress field, resulting in projection-like trajectories. The selected range ($t^* = 100–300$) represents a stable regime that yields anatomically consistent U-fiber formation and robust gyrus-to-gyrus connectivity, consistent with *in vivo* imaging observations of SAF organization.

In our recent study(*130*), we presented the values and ranges of previously reported elongation rates that depend on stretch or force, along with the calculated stress-induced fiber elongation rate, denoted as $a$. The axonal growth rate, $G^{\mathrm{axn}}$, exhibits significant variability, ranging from $0.2$ to $1.8\ \mathrm{mm\,d^{-1}}$ and the induced elongation rate spans from $0.0004$ to $0.31$ $\mathrm{mm\,Pa^{-1}\,d^{-1}}$. For our model, we assume fixed axonal growth rates of $G^{\mathrm{axn}} = 0.6$, $0.8$, $1.0$, and $1.2\ \mathrm{mm\,d^{-1}}$ and set a stress-induced elongation rate of $a = 0.015\ \mathrm{mm\,Pa^{-1}\,d^{-1}}$. Parameter $a$ was selected based on literature values and a sensitivity analysis presented in Fig. S7. In regions far from the cortex where stress is minimal or absent, fibers initiate growth at a steady rate of $G^{\mathrm{axn}}$. Here, fibers exhibit a less directed, more random growth trajectory, consistent with previous observations(*113*, *132*). However, in areas where cortical folding generates mechanical stress, axons reorient to align with the MTPS direction. This alignment enhances the growth rate to $G^{\mathrm{axn}} + 0.015\sigma\ \mathrm{mm\,d^{-1}}$, where $\sigma$ is the local stress. To accurately model the mechanical behavior of fibers during growth and deformation, we implement a nonlinear hyperelastic material model, which is a suitable first-order approximation for capturing the elastic and stretch-responsive properties of axons. This approach, which aligns with our micromechanical model(*96*) or other models in the literature(*97*), allows the fibers in our simulation to undergo stretching, elongation, or compression as the surrounding brain tissue deforms under the influence of folding processes.

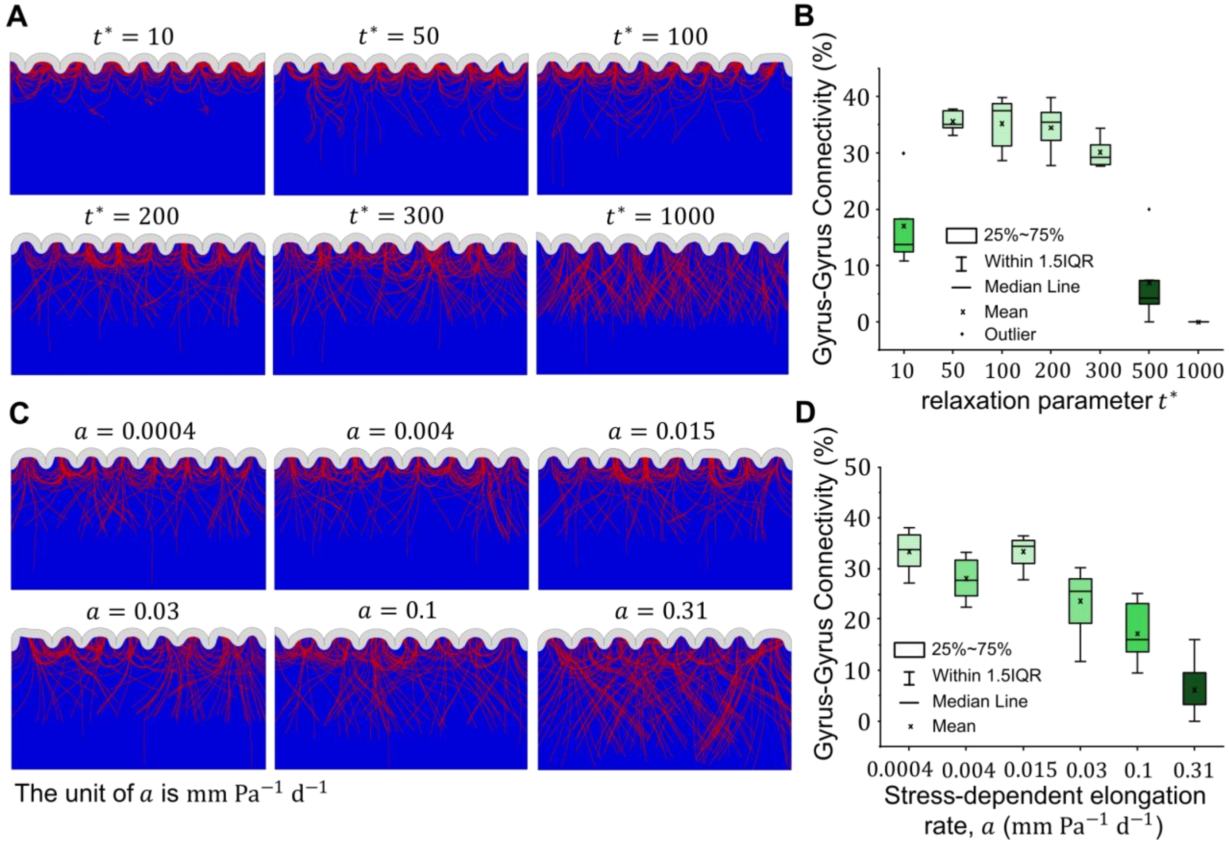

**Fig. S7. Sensitivity of fiber organization and gyrus-to-gyrus connectivity to the stress-dependent elongation rate $a$ and reorientation timescale.** (**A**) Representative simulations of cortical folding and axonal fiber development for increasing values of the relaxation parameter $t^*$ (10, 50,100, 200, 300, 500, and 1000). For very small $t^*$ (e.g., 10), fibers reorient too rapidly, leading to abrupt and non-physical trajectories, including local looping at fiber tips. At $t^* = 50$, fibers remain overly confined within individual gyri and fail to consistently connect adjacent gyri. In the intermediate range ($t^* = 100\text{-}300$), fibers exhibit smooth, gradual reorientation and form realistic U-shaped SAFs bridging neighboring gyri. For larger values ($t^* \geq 500$), reorientation becomes too slow relative to the evolving stress field, and fibers tend to follow elongated, projection-like trajectories; at $t^* = 1000$, SAF formation is largely suppressed. (**B**) Gyrus-to-gyrus connectivity quantified for all values of $t^*$ (each case repeated 10 times). Connectivity remains relatively stable for $t^* = 100\text{-}300$, indicating a robust regime of SAF formation. For larger values ($t^* \geq 300$), connectivity decreases significantly, consistent with reduced reorientation and increased projection-like fiber behavior. (**C**) Representative simulations of cortical folding and axonal fiber development for increasing values of the stress-dependent elongation rate $a$, spanning 0.0004 to 0.310. Each case shows the resulting fiber trajectories within the folded geometry. For lower values of $a$, fibers exhibit gradual reorientation and form localized U-shaped SAFs. As $a$ increases, fibers elongate more rapidly in response to stress, leading to reduced reorientation and more projection-like trajectories. (**D**) Gyrus-to-gyrus connectivity quantified across all values of $a$ (each case repeated 10 times). Connectivity remains relatively stable for $a \leq 0.015$, indicating low sensitivity in this regime. For larger values ($a > 0.03$), both the number of SAFs and gyrus-to-gyrus connectivity decrease significantly. At high $a$ (e.g., $a = 0.31$), rapid stress-dependent elongation dominates, limiting the ability of fibers to reorient toward the local MTPS, resulting in reduced SAF formation. The box plots represent the interquartile range (IQR), extending from the 25th to the 75th percentile. The mean is depicted as a white dot, while the median is shown as a black line within the box. The whiskers indicate the data range within 1.5 times the IQR. The parameters used in this figure are as follows: $\mu_\mathrm{f}/\mu_\mathrm{s} = 2$, $\mu_\mathrm{c}/\mu_\mathrm{s} = 2$, $G^\mathrm{axn} = 0.6\ \mathrm{mm\ d^{-1}}$, where $\mu_\mathrm{f}$, $\mu_\mathrm{s}$, and $\mu_\mathrm{c}$ are the shear moduli of the fiber, subplate, and cortex, respectively, and $G^\mathrm{axn}$ is the axon growth rate.

In our simulations, the shear modulus ratio of the cortex to the subplate was initially set at $2$. The shear modulus ratio between axons and white matter ECM has been reported within a broad range, based on studies primarily focused on mature, highly myelinated axons(*98*, *133*). However, for a developing brain with minimal myelination, it is expected that this ratio is lower, as myelination significantly contributes to white matter stiffness. Indeed, white matter stiffness is known to increase as myelin content rises(*91*). For our baseline model, we set the axon-to-subplate shear modulus ratio to $2$, with plans to investigate how varying this ratio affects axonal growth and the cortical folding process. Table 1 provides a comprehensive list of the parameters and associated values used in our simulations.

Our supplemental analyses showed that increasing the number of fibers beyond approximately 150 has minimal impact on the resulting fiber density in both gyri and sulci after folding, as well as on regional connectivity. This stabilization is demonstrated in Fig. S8A, where the statistical outcomes become consistent and largely independent of fiber count beyond this threshold. Additionally, we performed a sensitivity analysis to assess the influence of fiber initiation positions. As shown in Fig. S8B, averaging the results over 10 trials per parameter set yields reliable and reproducible statistical outcomes, minimizing variability due to random initialization.

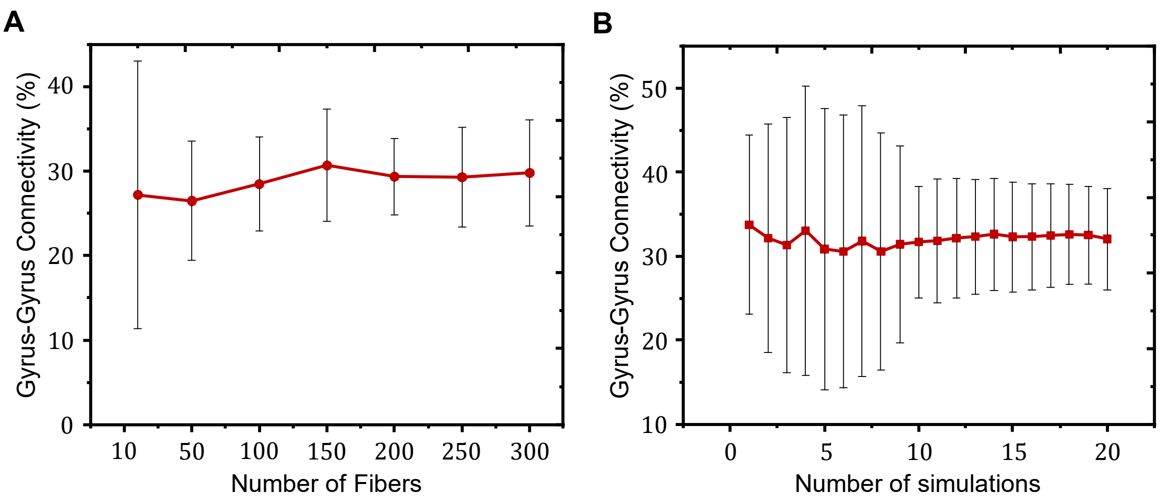


**Fig. S8. Effect of fiber number and number of simulations on gyrus-to-gyrus connectivity.** (**A**) The number of fibers does not impact the percentage of gyrus-to-gyrus streamlines beyond 150. (**B**) Relationship between the number of simulations on gyrus-to-gyrus connectivity. Results demonstrate stability in percentage of gyrus-to-gyrus streamlines beyond a threshold of 10 simulations. The used parameters in this figure are: $\mu_f/\mu_s = 2$, $\mu_c/\mu_s = 2$, $a = 0.015\ \mathrm{mm\ Pa^{-1}\ d^{-1}}$, $G^{\mathrm{axn}} = 0.6\ \mathrm{mm\ d^{-1}}$, where $\mu_f$, $\mu_s$, and $\mu_c$ are the shear moduli of the fiber, subplate, and cortex, respectively, $a$ is the stress-dependent elongation rate, and $G^{\mathrm{axn}}$ is the axon growth rate. Data are represented as mean values +/-SD (Standard Deviation).

### Brain imaging data and processing

In this study, we used grayordinate T1/2-weighted (T1/2w) MRI data and dMRI data from the Human Connectome Project (HCP) 3T dataset, provided by the Washington University-Minnesota consortium(*117*). The age range of the participants was between 22 and 35 years, with a gender distribution that was approximately balanced. For the purposes of our analysis, we randomly selected a cohort of 14 subjects from the HCP 3T dataset. All participants provided written informed consent, and the study received approval from the relevant institutional review boards. The parameters of T1w data of the HCP 3T dataset are as follows: TR=2400 ms, TE=2.14 ms, flip-angle=8 deg, image matrix=260×260, FOV=224 mm$^2$, and resolution=0.7×0.7×0.7 mm$^3$. T2w: TR=3200 ms, TE=565 ms, image-matrix=260×311×260, FOV=224 mm$^2$, and resolution=0.7×0.7×0.7 mm$^3$. dMRI: TR = 5520 ms, TE = 89.5 ms, refocusing flip-angle = 160$^0$, flip-angle = 78$^0$, FOV = 210×180 mm$^2$, matrix = 168×144, resolution = 1.25×1.25×1.25 mm$^3$, 1.25 mm isotropic voxels, echo-spacing = 0.78 ms, BW = 1,488 Hz/Px, three shells of b-values = 1000, 2000, and 3000 s/mm$^2$, with three different gradient tables, each of which includes approximately 90 diffusion weighting directions.

*Diffusion MRI preprocessing and short association fibers extraction.* The overall workflow for the imaging data extraction is summarized in Fig. 7. Diffusion-weighted imaging (DWI) data preprocessing was conducted using established pipelines in MRtrix3.0(*118*) and ANTs(*119*). Thermal noise was suppressed using a principal component analysis (PCA)-based denoising approach(*134*), which enhances the signal-to-noise ratio and facilitates more robust estimation of diffusion metrics. Motion and eddy current distortions were addressed using a combination of rigid-body alignment and tensor-based registration, with subsequent co-registration to high-resolution T1-weighted anatomical images via symmetric diffeomorphic transformations(*119*). A five-tissue-type (5TT) classification map was generated to distinguish cortical gray matter (GM), subcortical GM, white matter (WM), cerebrospinal fluid (CSF), and pathological tissue(*135*). This segmentation leveraged probabilistic tissue models derived from T1-weighted anatomical images and was critical for enforcing anatomically constrained tractography (ACT) to ensure biologically plausible streamline trajectories(*136*).

Multi-shell, multi-tissue (MSMT) response functions were estimated to model signal contributions from WM, GM, and CSF(*137*). Fiber orientation distributions (FODs) were reconstructed using constrained spherical deconvolution (CSD), which resolves crossing fiber orientations by deconvolving the measured diffusion signal with a reference response function(*138*). FOD amplitudes were normalized to ensure tissue-specific signal contributions remained physiologically interpretable(*137*).

Whole-brain probabilistic tractography was performed using a second-order integration algorithm over FODs, dynamically seeding streamlines within WM voxels. Streamlines were constrained by the 5TT map to terminate at GM-WM interfaces(*136*). To isolate short association fibers (SAFs), a maximum streamline length threshold of 40 mm was applied(*1*). A custom computational pipeline was developed to filter SAFs based on strict anatomical criteria. Streamlines were retained only if both endpoints resided within the neocortical GM of the same hemisphere, with intermediate segments traversing WM. Streamlines terminating in subcortical structures, crossing hemispheres, or exceeding the 40 mm length threshold were excluded. The WM-GM interface was reconstructed using a surface-based morphometry pipeline(*139*). SAF endpoints are mapped to cortical regions to compute short-range connectivity matrices representing gyrus–gyrus interactions.
